# CUTS RNA Biosensor for the Real-Time Detection of TDP-43 Loss-of-Function

**DOI:** 10.1101/2024.07.12.603231

**Authors:** Longxin Xie, Jessica Merjane, Cristian A Bergmann, Jiazhen Xu, Shruthi Balasubramaniyan, Bryan Hurtle, Charleen T Chu, Christopher J Donnelly

**Author notes:** Authors share equal contributions (Longxin Xie, Jessica Merjane, Cristian A Bergmann).

## Abstract

Mounting evidence implicates TDP-43 dysfunction and the accumulation of pathological cryptic exons across multiple neurodegenerative diseases, underscoring the need for accessible tools to detect and quantify TDP-43 loss-of-function (LOF). These tools are crucial for assessing potential disease contributors and exploring therapeutic candidates in TDP-43 proteinopathies. Here, we develop a sensitive and accurate real-time sensor for TDP-43 LOF: the CUTS (CFTR UNC13A TDP-43 Loss-of-Function) system. This system combines UG-rich sequences and previously reported cryptic exons regulated by TDP-43 with a reporter, enabling the tracking of TDP-43 LOF through live microscopy and RNA/protein-based assays. We show that CUTS effectively detects TDP-43 loss of function arising from mislocalization, impaired RNA binding, and pathological aggregation. Our results show the sensitivity and accuracy of the CUTS system in detecting and quantifying TDP-43 LOF, opening avenues to explore unknown TDP-43 interactions that regulate its function. In addition, by replacing the fluorescent tag in the CUTS system with the coding sequence for TDP-43, we show significant recovery of its function under TDP-43 LOF conditions, highlighting the potential utility of CUTS for self-regulating gene therapy applications. In summary, CUTS represents a platform for evaluating TDP-43 LOF in real-time and gene-replacement therapies in neurodegenerative diseases associated with TDP-43 dysfunction.

**Highlights:**

- CUTS is a cryptic exon RNA biosensor enabling real-time detection of TDP-43 loss of splicing function.
- CUTS exhibits a linear relationship with a reduction in TDP-43 protein.
- CUTS can deliver an autoregulated gene payload in response to TDP-43 loss-of-function.
- TDP-43 homotypic phase transitions and cell stress induce loss of splicing function detected via CUTS.

## Introduction

Amyotrophic lateral sclerosis (ALS) is a progressive and fatal neurodegenerative disease (NND) characterized by a persistent degeneration of the neurons of the spinal cord and motor cortex (Feldman et al., 2022). The dysregulation of the RNA binding protein (RBP), TAR DNA-binding protein 43 (TDP-43) is a hallmark pathobiology observed in ∼97% of all ALS patients (Neumann et al., 2006), ∼45% of Frontotemporal lobar degeneration (FTLD) patients (Ling et al., 2013; Neumann et al., 2006), and 40%-60% of Limbic Associated TDP-43 Encephalopathy (LATE) patients (McKee et al., 2010; Nelson et al., 2019; Tremblay et al., 2011). Under physiological conditions, TDP-43 orchestrates many cellular processes critical for neuronal health and homeostasis, including regulating RNA metabolism, splicing, and stress response pathways. However, in disease, TDP-43 is depleted from the nucleus and mislocalizes to the cytoplasmic compartment, losing the ability to perform its canonical splicing function and may transition into insoluble aggregates (Hurtle et al., 2023; Mann and Donnelly, 2021).

Recent efforts increasingly focus on the implications of TDP-43’s loss of splicing function and its consequence on disease onset and progression (Brown et al., 2022; Buratti and Baralle, 2001; Klim et al., 2019; Ling et al., 2015; Ma et al., 2022; Melamed et al., 2019). Physiologically, TDP-43 selectively binds to specific UG-rich sequences within pre-mRNA transcripts, providing precise control over the alternative splicing of a subset of conserved targets, thereby modulating their gene expression and cellular function (Kuo et al., 2009; Lukavsky et al., 2013). This regulatory function is key in repressing the incorporation of TDP-43-mediated aberrant ‘cryptic exons’ (CE), non-conserved intronic regions whose inclusion has been linked to cellular toxicity and pathological consequences (Brown et al., 2022; Klim et al., 2019; Ling et al., 2015; Ma et al., 2022; Mehta et al., 2023; Melamed et al., 2019; Polymenidou et al., 2011). Noteworthy examples of transcripts susceptible to aberrant CEs in the absence of functional TDP-43 include *STMN2* (Klim et al., 2019; Melamed et al., 2019) and *UNC13A* (Brown et al., 2022; Ma et al., 2022), both pivotal in maintaining the integrity and physiological function of neurons, such as axon regeneration and motor neuron firing (Shin et al., 2014, 2012; Varoqueaux et al., 2002; Willemse et al., 2023). Therefore, their aberrant splicing patterns and loss-of-function (LOF) in disease highlights the critical role of TDP-43 in preserving neuronal health and function through the regulation of key neuronal health.

A well-established and currently available method to accurately monitor and quantify TDP-43’s splicing function has frequently relied on employing cryptic exon 9 inclusion in the human cystic fibrosis transmembrane conductance regulator (CFTR) transgene (Buratti et al. 2001; Ayala et al. 2006). However, this approach can only be used at the RNA level, limiting its functionality as a reporting system for high-throughput screens or *in vivo* applications. Furthermore, endogenous cryptic exons exhibit variable responses to TDP-43 loss, may be cell type-specific, and are still being defined. To address this, we designed a screening tool, the ‘CFTR-UNC13A TDP-43 loss-of-function Sensor’ (CUTS) system, engineered to detect TDP-43 LOF and output a proportional and quantifiable signal using GFP or other reporters. By combining previously described TDP-43 binding targets with small UG stretches, we significantly improved the sensitivity measurable in real time using standard assays. Functional TDP-43 promotes the splicing of CUTS CE, resulting in a frameshift and early stop codon that prevents GFP expression. TDP-43 LOF activates the CUTS biosensor by promoting the inclusion of the CE, allowing GFP to be expressed. Here, we highlight the CUTS system sensitivity in response to TDP-43 loss. The CUTS design enables GFP expression to be directly proportional to TDP-43 LOF, allowing the detection of changes in TDP-43 function even when alterations cannot be measured directly by traditional methods like qPCR, western blot, or immunofluorescent imaging.

In this study, we highlight the utility and sensitivity of the CUTS biosensor in non-neuronal and neuronal cells. We also show that aberrant TDP-43 phase transitions or mislocalization disrupt TDP-43 splicing function by expressing well-established RNA-binding-deficient and cytoplasmic TDP-43 constructs. We also replaced the GFP cassette in the CUTS biosensor with the wild-type *TARDBP* coding sequence (CUTS-TDP43), demonstrating that CUTS can regulate a gene payload expression in response to TDP-43 loss-of-function. The CUTS biosensor provides a sensitive and reliable means to dynamically monitor TDP-43 function and can be applied for therapeutic screening, assessment of disease-associated dysfunction, and delivery of genetic payloads in response to TDP-43 loss.

## Results

### CUTS TDP-43 LOF sensor design utilizing known Cryptic Exons

To improve the detection of TDP-43 LOF, we designed a novel TDP-43 LOF sensor (TS) using previously reported genes known to undergo TDP-43-regulated splicing (*UNC13A* and *CFTR*). The TS cassettes are constructed with a constitutively expressed mCherry, followed by a TDP-43-regulated CE and a GFP linked to a 3x nuclear localization signal (NLS), each separated by a T2A self-cleavage sequence (Figure 1A). We positioned the GFP reporter outside the mCherry open reading frame (ORF), introducing an early stop codon upstream of the GFP (Figure 1A). This strategic design achieved two key outcomes: (1) Under physiological TDP-43 levels, the binding of TDP-43 to the CE and UG-rich sequence should promote complete intronic splicing, maintaining the in-frame stop codon upstream of GFP, resulting in only mCherry expression. (2) TDP-43 loss and/or failure to bind the CE sites will result in CE retention and a subsequent frameshift that bypasses the stop codon, allowing translation to proceed into the GFP coding sequence, leading to co-expression of mCherry and GFP. This system will allow a systematic screen of whether any TDP-43 perturbation is causing LOF, which should be proportional to GFP signal output.

**Figure 1:**
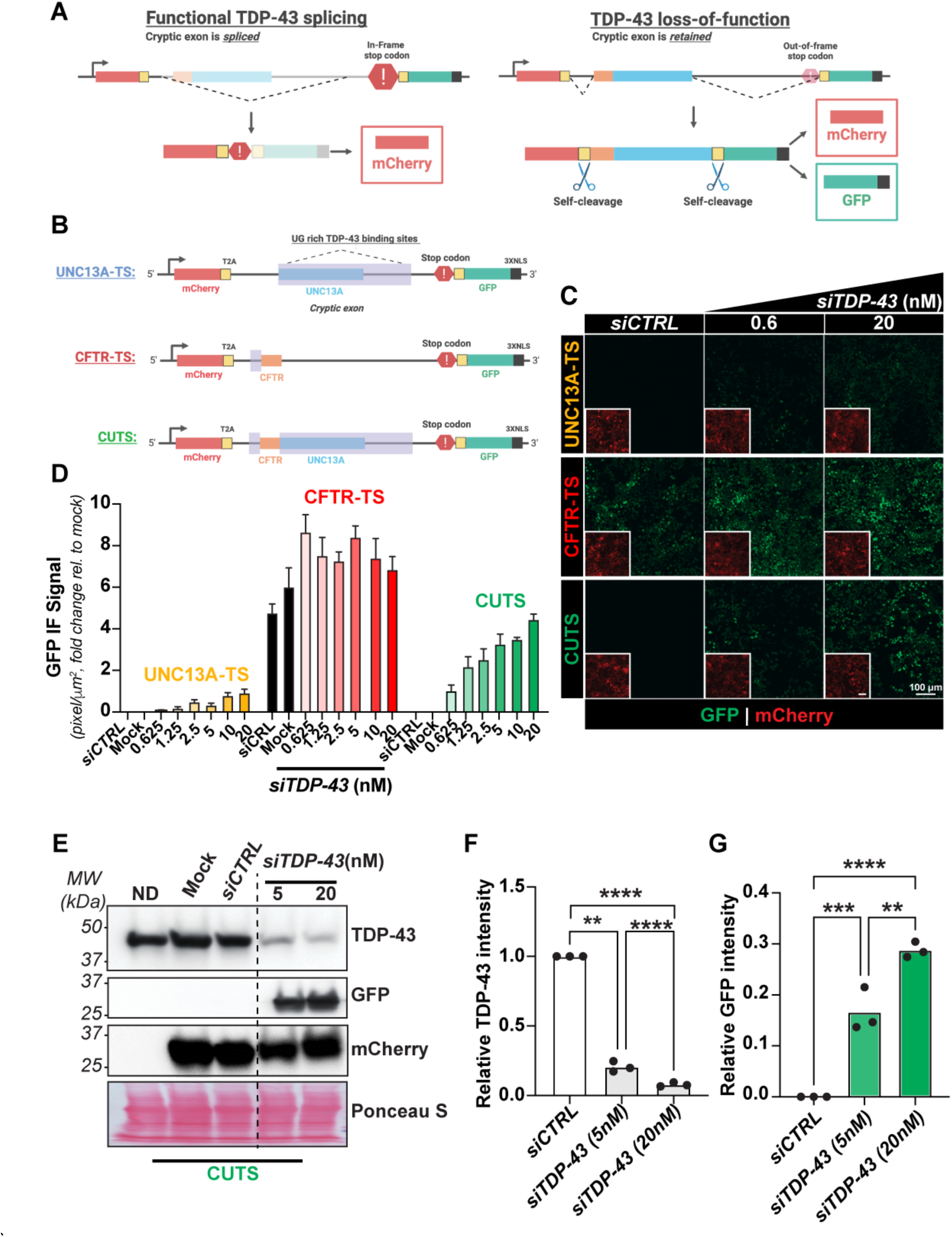
Design and functional comparison of TDP-43 loss-of-function reporter systems. Comparison of stable polyclonal HEK cells expressing UNC13A-TS, CFTR-TS, or CUTS following treatment with siRNA control (siCTRL) (20nM) or TDP-43 (siTDP-43) (0.6nM - 20nM). Cells were reverse-transfected with siRNA treatment in complete media supplemented with doxycycline (1000 ng/mL). As a negative control, we use ND: No doxycycline. After 72 h, cells were analyzed by live imaging and protein lysate was harvested for western blot analysis. **(A)** Schematic of the TDP-43 loss-of-function Sensor (TS) system design. **(B)** Overview of the UNC13A-TS, CFTR-TS, and CUTS cryptic exon cassette design. **(C)** Representative live imaging of TS comparison (10X) nuclear GFP indicates TDP-43 LOF. **(D)** Mean intensity quantification of GFP signal intensity as shown in (C). **(E)** Western blot analysis of CUTS immunoblotting for TDP-43 and GFP proteins. **(F-G)** Relative pixel quantification of TDP-43 and GFP band normalized to total protein (Ponceau S) from (E). Statistical significance was determined by one-way ANOVA and Tukey’s multiple comparison test (*, p < 0.05; **, p< 0.01; ***, p < 0.001; ****, p < 0.0001). Green = GFP signal; red = mCherry signal. Scale bar = 100 µm. N=3 biological replicates.

**Supplementary Figure 1:**
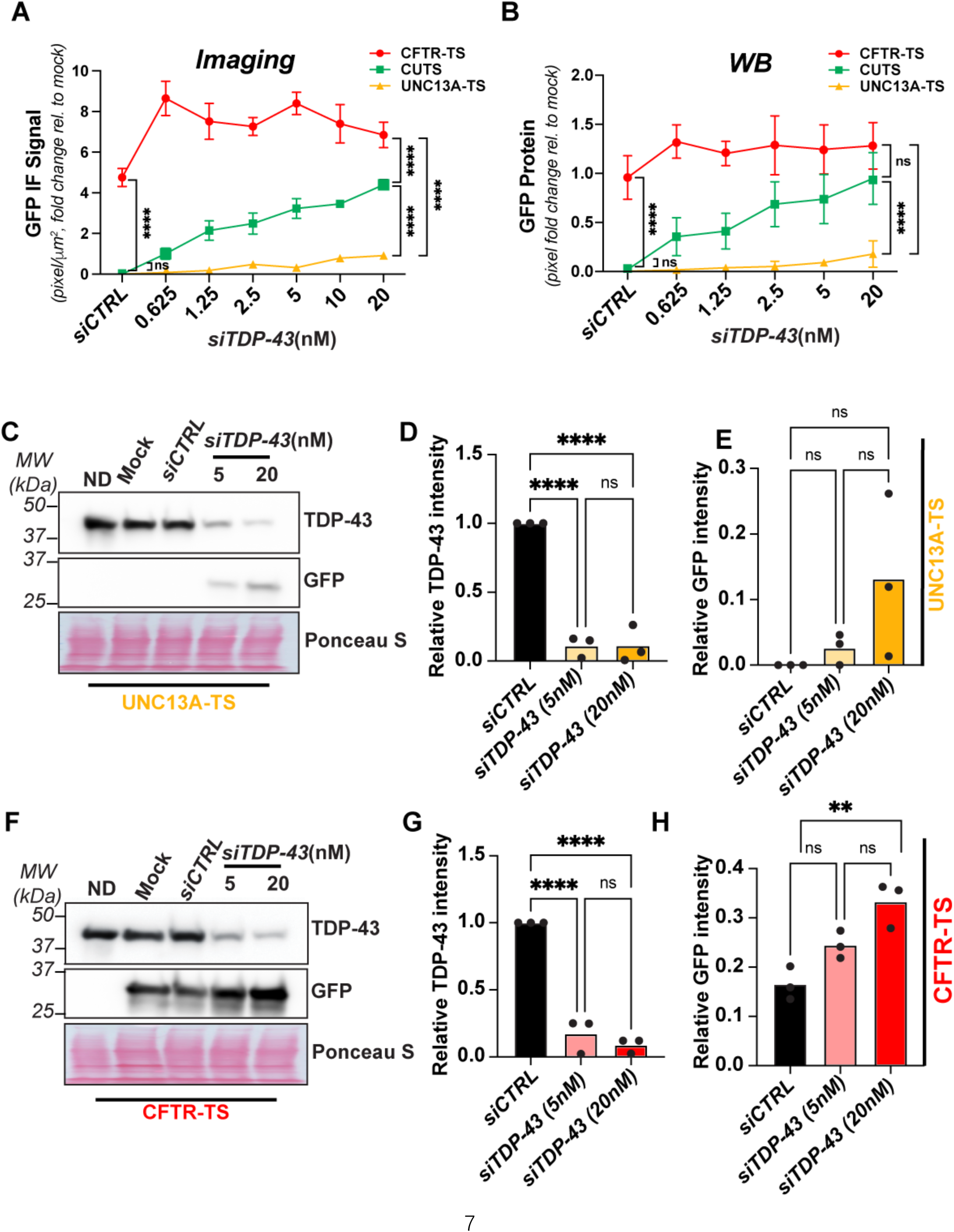
Differential reporter sensitivity to TDP-43 depletion. Comparison of stable polyclonal HEK cells expressing UNC13A-TS, CFTR-TS, or CUTS following treatment with siRNA control (siCTRL) (20nM) or TDP-43 (siTDP-43) (0.6nM - 20nM). Cells were reverse-transfected with siRNA treatment in complete media supplemented with doxycycline (1000 ng/mL). After 72 h, cells were analyzed by live imaging and protein lysate was harvested for western blot analysis. (**A)** Mean intensity quantification of GFP signal intensity from live imaging, presented as fold change from mock. **(B)** Relative pixel quantification of GFP normalized to total protein (Ponceau S), presented as fold change from mock. **(C-E)** WB analysis of UNC13A-TS, developing again TDP-43 and GFP proteins, and pixel quantification of TDP-43 and GFP intensity from bands show in (C) normalized to total protein (Ponceau S). **(F-H)** WB analysis of CFTR-TS, developing again TDP-43 and GFP proteins, and pixel quantification of TDP-43 and GFP intensity from bands show in (F) normalized to total protein (Ponceau S). Statistical significance was determined by two-way ANOVA and Tukey’s multiple comparison test (*, p < 0.05; **, p < 0.01; ***, p < 0.001; ****, p < 0.0001). N=3 biological replicates.

### CUTS is more precise than CFTR and UNC13A as a TDP-43 LOF biosensor

To assess the functionality of TS, we engineered UNC13A-TS, CFTR-TS, and CUTS, which utilized the TDP-43-regulated CEs from UNC13A, CFTR, and a combined construct termed CUTS (CFTR-UNC13A TS), integrating both CE sequences (Figure 1B). Additional base modifications were incorporated into the cassette designs to prevent unexpected stop codons within the CE regions (see Table S1 for full sequence details). Each construct was coupled with a Tet3g promoter, cloned into a Piggybac vector, and stably expressed in HEK293 cells. To evaluate the accuracy and reliability of the three TS constructs, we performed a TDP-43 LOF assay using increasing *siRNA* concentrations, followed by live confocal imaging and western blot (WB) analysis (Figure 1C-G and S1A-H). Across all cell lines, we observed constitutive mCherry expression and a trend toward increased GFP signals with higher *siTDP-43* concentrations. The UNC13A-TS construct demonstrated no detectable GFP expression under control conditions in both imaging-based and WB analyses (Figure 1 C-D, S1A-E). However, the UNC13A-TS showed limited sensitivity, with only a modest GFP signal (17% of CUTS) detected via imaging under the highest dose of siTDP-43 treatment assessed (20nM; Figure 1D). In contrast, the CFTR-TS construct exhibited a notably high baseline, with detectable GFP leakage in control groups without TDP-43 loss (Figure 1C-D, S1A-B, F-H). Interestingly, cells expressing the CUTS construct exhibited a synergistic effect from both the CFTR and UNC13A CE sequences, achieving high sensitivity evidenced by a clear dose-responsive GFP expression with increasing siTDP-43 concentrations, while maintaining high accuracy with minimal leakage via imaging and GFP immunoblotting (Figure 1C-G). Given the promising accuracy of the CUTS sensor, we proceeded to further characterize the CUTS RNA biosensor.

### CUTS demonstrates ultra-sensitivity in detecting low-level TDP-43 loss-of-function

To challenge the stability and sensitivity of CUTS, we next conducted an ultra-low dose TDP-43 *siRNA* transfection, ranging from 38-1200 pM. Immunofluorescence staining (IF) revealed a consistent increase in both GFP intensity and GFP-positive cell ratios in CUTS-expressing cells in response to elevated *siTDP-43*, with minimal baseline expression observed (Figure 2A-B). We confirmed the ultra-sensitivity and accuracy of CUTS using WB analysis, demonstrating measurable GFP even at the lowest doses of *siTDP-43* assessed (Figure 1C-E). While changes in TDP-43 levels were undetectable by WB at *siTDP-43* doses of 37.5-75 pM (measured as 2% TDP-43 KD by WB, see Figure S2A), the CUTS system demonstrated a 7 to 55-fold increase in GFP expression at these doses compared to baseline (Figure S2A). This increase in GFP expression persisted to the highest siTDP-43 dose (1,200 pM; 98% TDP-43 KD; Figure S2A), resulting in a 118,224-fold increase in GFP compared to baseline. Pearson’s correlation analysis confirmed a highly significant relationship between GFP expression and siTDP-43 dose (*p*=0.0011), while the correlation between measurable TDP-43 and siTDP-43 concentration was less significant (*p*=0.0429) (Figure 2D-E). Linear regression analysis between the logarithmic GFP fold increase and siRNA doses demonstrated an exceptional linear relationship (R^2^=0.9998) (Figure 2E). These results indicate that GFP expression produced by CUTS is a more sensitive method for detecting TDP-43 KD (and therefore LOF) than TDP-43 WB detection. The linear relationship between GFP and siTDP-43 dose also demonstrates the potential of CUTS to be used as a predictive model for TDP-43 LOF.

**Figure 2:**
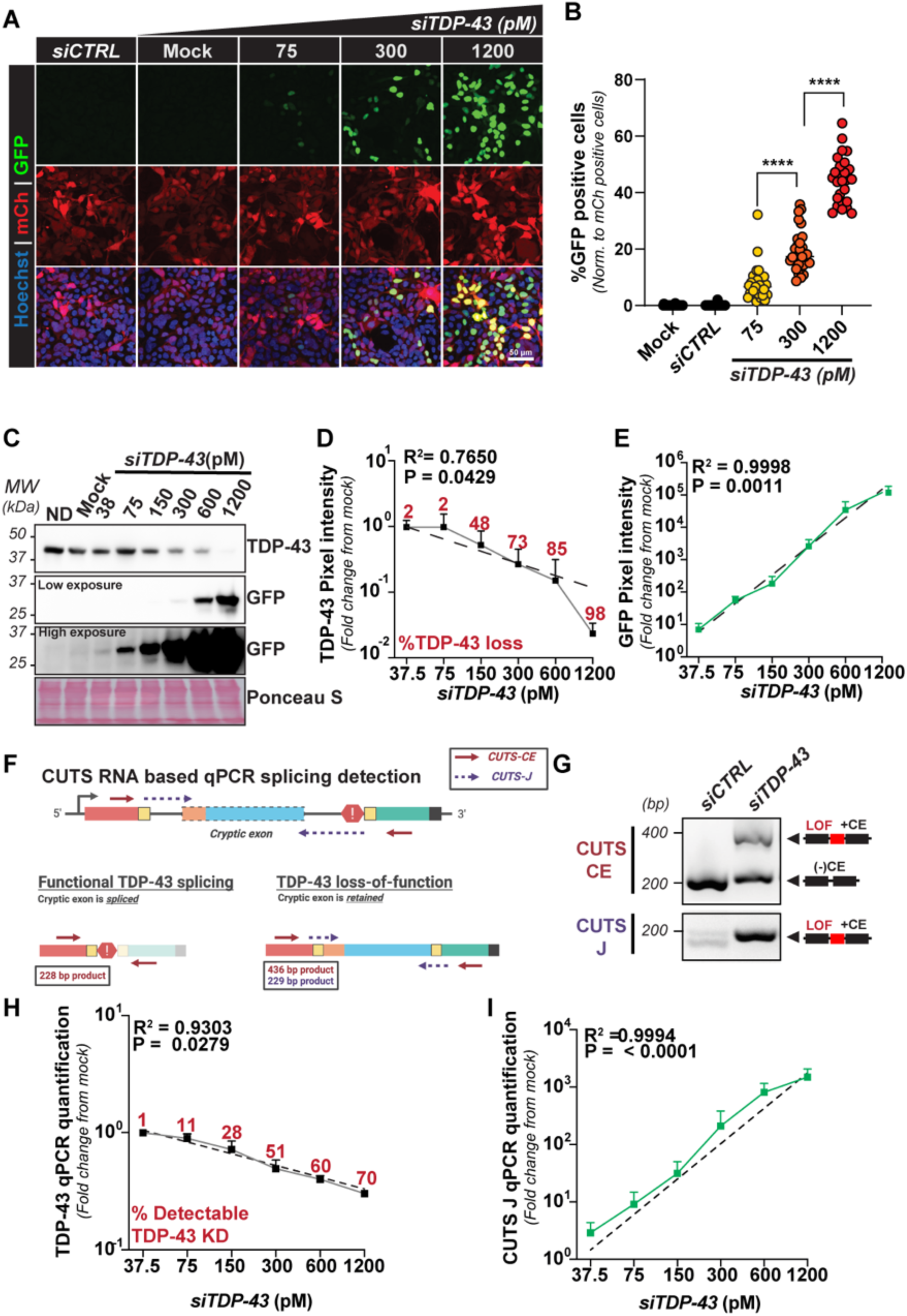
CUTS exhibits linear, dose-responsive activation following low-level TDP-43 knockdown. Low-dose siRNA TDP-43 (siTDP-43) treatment was performed in stable polyclonal HEK cells expressing CUTS. CUTS-expressing cells were reverse transfected with siRNA control (siCTRL) or siTDP-43 in a dose-response curve (38 to 1200pM) in doxycycline supplemented media (1000ng/ml) for 72hr. **(A)** Representative immunofluorescence images of CUTS-expressing HEK cells under low doses of siRNA TDP-43 treatment. (60X). **(B)** Percentage of GFP-positive cells normalized to the number of CUTS-expressing mCherry-positive cells. **(C)** Western blot of TDP-43 and GFP proteins from HEK cell lysate expressing CUTS under low doses of siRNA TDP-43. Ponceau S is shown as a loading control. **(D)** Pixel intensity quantification of the TDP-43 and **(E)** GFP bands shown in (C), presented as fold-change from the mock-treated sample. **(F)** Schematic showing the position of qPCR primers, developed to detect CUTS cryptic exon inclusion (referred to as ‘CUTS-CE’ and ‘CUTS-J’). **(G)** Representative agarose gel showing qPCR product from melting curve detecting CUTS cryptic exon inclusion using the primers shown in (F). **(H-I)** qPCR quantification of TDP-43 and CUTS cryptic exon J from the siTDP-43 dose curve presented as fold-change from the mock-treated sample. Linear regression analysis shown in (D), (E), (H) and (I) was performed on Log values. Fitting method = least squares regression. (****, p < 0.0001). N=3 biological replicates. Green = GFP; red = mCherry. Scale bar = 50 µm. N=3 biological replicates.

We assessed CUTS’ sensitivity at the transcript level using the *siTDP-43* ultra-low dose curve and RT-qPCR assessment in Figure 2F-I. To determine the relative amount of CUTS’ CE retention, we designed primers targeting either the entire transcript or the specific junction sites of the CE (Figure 2F-G). The RT-qPCR quantification demonstrated increased sensitivity at detecting changes in TDP-43 levels compared to WB analysis, with the capability of detecting changes in TDP-43 between each siTDP-43 dose (1 – 70% TDP-43 KD, see Figure 2H and Figure S2B) in a linear manner (R^2^ = 0.9303). Using CUTS detection, we observed a clear linear logarithmic relationship between the amount of CUTS CE-retention and the increasing *siTDP-43* doses (R^2^=0.9994) (Figure 2I and Figure 2SB). Even at 1% KD in TDP-43, the CUTS system detected a 3-fold increase in CE retention compared to baseline, which increased to a 1,488-fold increase at 70% TDP-43 KD (Figure 2I and Figure 2SB). As with our WB analysis (Figure 2D), there was a significant correlation between GFP expression and TDP-43 and the siTDP-43 dose (p < 0.0001 and p = 0.0279, respectively). Thus, this data suggests that CUTS is a reliable tool to quantify TDP-43’s function across an extensive functional range, highlighting its utility as a TDP-43 LOF biosensor.

Additionally, CUTS demonstrated ultra-sensitivity under low-level TDP-43 KD, beyond the detection limit of both WB and RT-qPCR. We next sought to compare the ultra-sensitive detection capacity of the CUTS sensor with the physiological response of CEs inclusion on genes regulated by TDP-43 (Figure S2E). Using low concentrations of siTDP-43, we were able to recapitulate CUTS sensitivity to detect CE inclusion at low-dose siRNA (75 pM) by RT-PCR. Interestingly, LPR8, ARHGAP32, and ACBD3 CEs displayed a similar pattern, showing progressive CE inclusion with increasing *siTDP-43* doses, starting at 75 pM. However, HDGFL2 showed higher sensitivity to TDP-43 LOF, with detectable exon inclusion even at 38 pM (Figure 2SE). Together, these results establish CUTS as a relevant tool to evaluate sensitively and quantitatively minimal reductions in TDP-43 levels and accurately report its LOF.

**Supplementary Figure 2:**
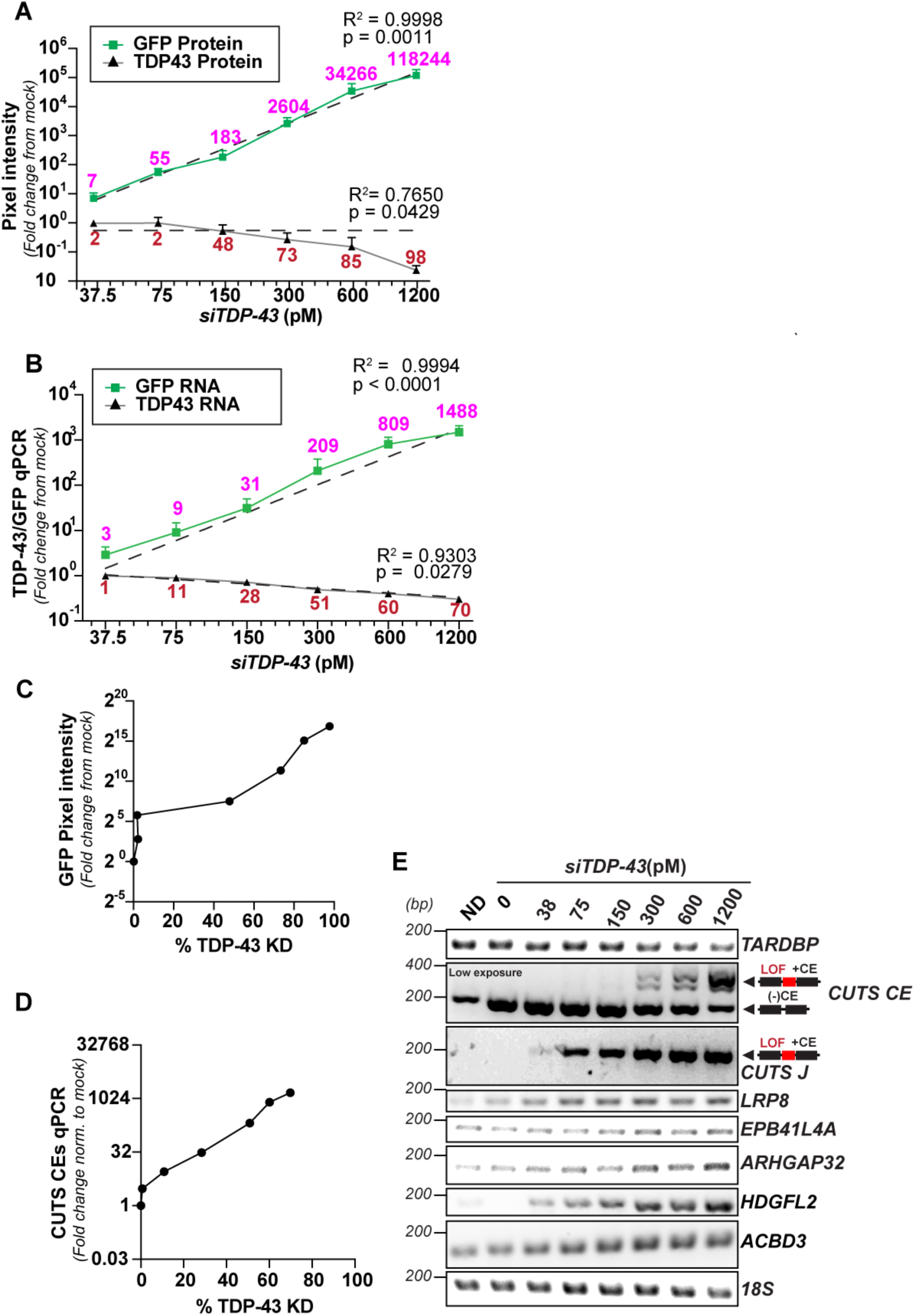
CUTS signal scales with dose-dependent TDP-43 knockdown. Low-dose siTDP-43 treatment was performed in stable polyclonal HEK cells expressing CUTS. CUTS-expressing cells were reverse transfected with siCTRL or siTDP43 in a dose-response curve (38 - 1200 pM) in doxycycline supplemented media (1000ng/ml) for 72 h. **(A)** Pixel intensity quantification of the GFP and TDP-43 bands shown in (C), presented as fold-change from the mock-treated sample. **(B)** qPCR quantification of the siTDP-43 dose curve presented as fold-change from the mock-treated sample. Purple text above indicates the GFP fold change from the mock-treated sample. Red text above indicates the percentage of total detectable TDP-43 knockdown. Linear regression analysis shown in (D) and (G) was performed on Log values. Fitting method = least squares regression. **(C)** GFP pixel intensity from the Western blot shown in Figure 2C plotted against the percentage of TDP-43 knockdown detected under siTDP-43 treatment (38-1200 pM). **(D)** CUTS CE transcript quantification plotted against the percentage of TDP-43 knockdown detected under siTDP-43 treatment (38 - 1200 pM). **(E)** RT-PCR of endogenous cryptic exons and the CUTS biosensor following increasing doses of siTDP43 treatment.

### Pathological TDP-43 phase transitions or mislocalization activate the CUTS biosensor

In ALS/FTLD, the absolute TDP-43 level remains largely unaffected. Instead, TDP-43 undergoes pathological mislocalization and/or phase transitions likely due to a reduction in RNA binding, which reduces the functional cellular TDP-43. To evaluate whether these events contribute to TDP-43 LOF, we tested CUTS’s ability to detect TDP-43 LOF caused by TDP-43 mislocalization or aggregation via aberrant phase transitions. We transfected CUTS HEK293 cells with four tagged TDP-43 isoforms: (1) TDP-43^WT^, (2) TDP-43^cyto^, (3) TDP-43^5FL^, and (4) TDP-43^cyto^ ^5FL^. The TDP-43^cyto^ variants contain point mutations located within the nuclear localization signal (NLS) of TDP-43, resulting in cytoplasmic mislocalization (Mann et al., 2019). The 5FL form contains five phenylalanine-to-leucine mutations within the two RNA recognition motif (RRM) domains of TDP-43 that greatly impaired TDP-43’s RNA binding ability and were previously reported to form aggregated "anisomes" inside the nucleus (Cohen et al., 2015; Elden et al., 2010; Mann et al., 2019; Yu et al., 2021). The TDP-43^cyto^ ^5FL^ combines both modifications, leading to insoluble cytoplasmic inclusions (Elden et al., 2010; Keating et al., 2023; Lu et al., 2022; Mann et al., 2019; Yu et al., 2021). Excluding TDP-43^WT^, all three modified versions have been proven to sequestrate endogenous TDP-43 into mislocalized or aggregated inclusions (Keating et al., 2023). Therefore, our objective was to utilize CUTS to determine whether the expression of these aggregation-prone TDP-43 variants elicits endogenous TDP-43 LOF.

The introduction of TDP-43^cyto^, TDP-43^5FL^, and TDP-43^cyto^ ^5FL^ induced nuclear GFP signal when assessed by immunofluorescence analysis, while neither the tagged plasmid backbone nor TDP-43^WT^ caused any detectable GFP (Figure 3A). We confirmed the expression of exogenous and endogenous TDP-43 levels by WB and quantified the relative GFP level in each condition (Figure 3B-3C). As we have previously shown that CUTS demonstrates a proportional response to TDP-43’s LOF (Figure 2), we were able to directly interpret the relative ability of the different TDP-43 mutants to trigger TDP-43 LOF by comparison of their GFP levels. All three TDP-43 mutants’ expression triggered significant LOF compared to the control conditions, albeit at varying significance levels. The most modest LOF effect was achieved by TDP-43^cyto^, followed by TDP-43^cyto^ ^5FL^, and TDP-43^5FL^ (Figure 3C). Interestingly, although both TDP-43^5FL^ and TDP-43^cyto^ ^5FL^ caused significantly elevated LOF, the TDP-43^5FL^ mutant alone mediated greater LOF than when combined with the NLS mutations highlighting the potential role of nuclear homotypic TDP-43 interactions potentially contributing to TDP-43 LOF in disease absent its cytoplasmic mislocalization. To further validate the functionality of exogenous TDP-43, we transfected TDP-43^WT^ into a HeLa TDP-43 knock-out cell line expressing CUTS (Roczniak-Ferguson & Ferguson, 2019). We detected a significantly decreased GFP signal compared to the backbone or non-transfected controls, which confirmed the full splicing function of TDP-43^WT^ (Figure S3A-S3D).

**Figure 3:**
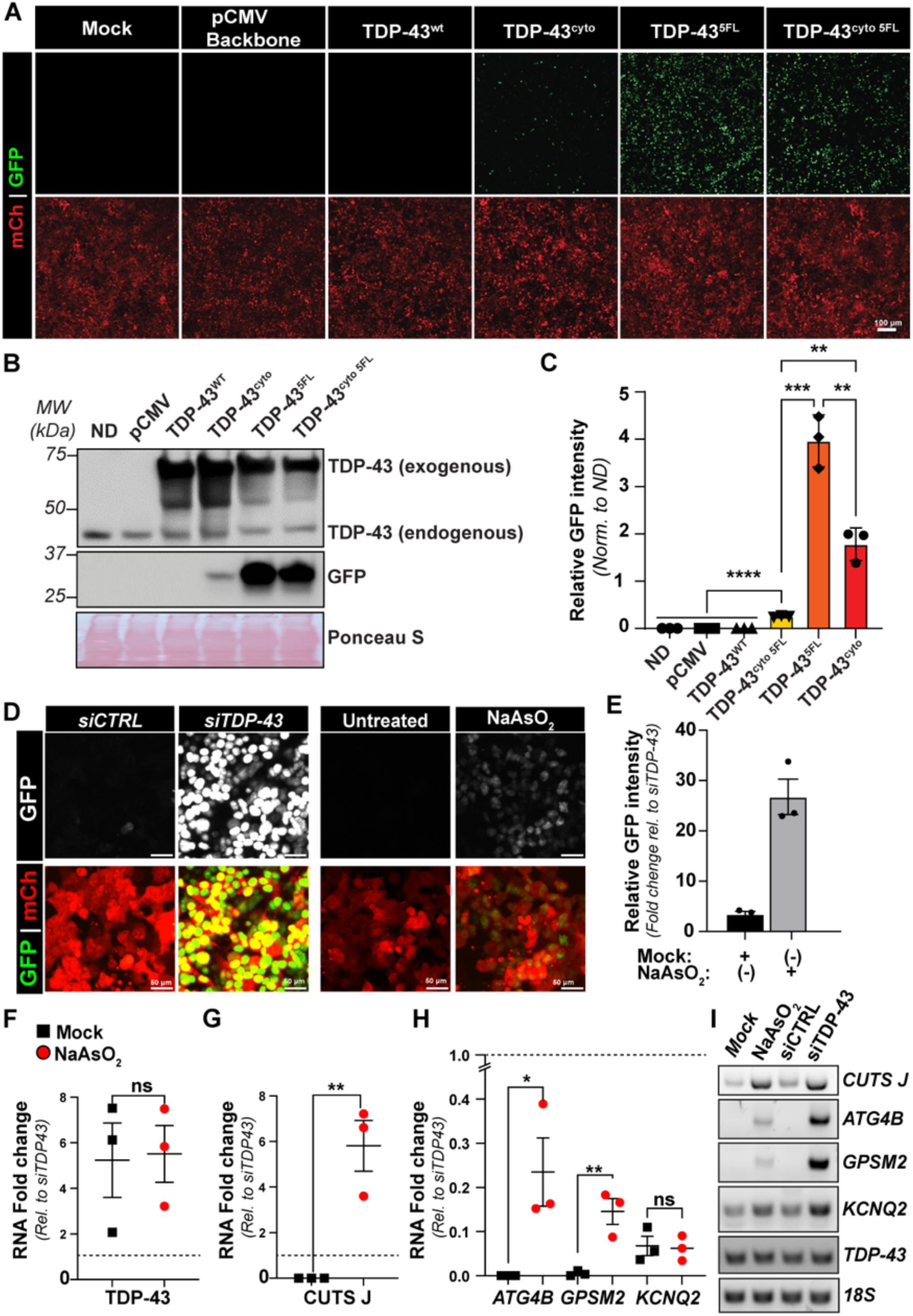
Mislocalization and aberrant phase transitions induce TDP-43 LOF. Stable HEK cells expressing CUTS were induced with doxycycline (1000 ng/mL) for 24 hours before transfection with the following plasmids: pCMV backbone, TDP-43^WT^, TDP-43^ΔNLS^, TDP-43^5FL^, TDP-43^ΔNLS^ ^5FL^, or non-transfected. Following transfection, plasmids were expressed for 72 h, followed by live imaging and protein analysis. **(A)** Live-imaging of CUTS HEK cells expressing WT or mutant TDP-43 gene cassettes. **(B)** Representative western blot of exogenous and endogenous GFP and TDP-43. Ponceau S is shown as a loading control. **(C)** Relative GFP pixel intensity quantification of the band is shown in (B). **(D–E)** Immunofluorescence of stable HEK cells expressing CUTS with siCTRL (20 nM), siTDP-43 (20 nM), mock, and NaAsO₂ (250 µM). **(E)** Immunofluorescence GFP signal quantification of (D) with 250 µM NaAsO₂ treatment relative to siTDP-43 (20 nM). **(F-G)** RT-qPCR from CUTS-expressing HEK cells treated with 250 µM NaAsO_2_ of TDP-43, CUTS-J transcript, and endogenous cryptic exons (ATGB4, GPSM2, and KCNQ2) relative to siRNA TDP-43 20 nM (dotted line) and normalized to 18S. **(I)** RT-PCR from Figure F-H. Expression values were normalized to 18S rRNA, and fold changes were calculated relative to the siTDP-43 (20 nM) condition. Black squares= Mock: Red circle= NaAsO₂. Statistical significance was determined by one-way ANOVA and Tukey’s multiple comparison test (*, p < 0.05; **. p < 0.01; ***. p < 0.001; ****, p < 0.0001). Green = GFP; red = mCherry. Scale bar = 100 µm. N = 3 biological replicates.

We next tested whether CUTS can detect TDP-43 LOF under oxidative stress induced by sodium arsenite (NaAsO₂), previously shown to disrupt TDP-43 splicing activity (Huang et al., 2024). CUTS showed mild activation, with detectable GFP signal and a significant increase in GFP intensity compared to mock treated cells (Figure 3D–E). While TDP-43 mRNA levels remained unchanged (Figure 3F), CUTS transcripts were markedly upregulated (Figure 3G). Consistent with this, endogenous cryptic exons ATG4B and GPSM2 were increased, whereas KCNQ2 was unaffected (Figure 3H-I)(Estades Ayuso et al., 2023; Joseph et al., 2025; Schmidt et al., 2019). These results highlight the sensitivity of CUTS in detecting subtle TDP-43 functional loss, which is often difficult to capture using an endogenous marker. Taken together, these data show the functional consequence caused by TDP-43’s mislocalization and/or aberrant phase transitions, demonstrating that aggregation-prone TDP-43 variants mediated direct LOF toxicity in addition to any gain-of-function (GOF) toxic events. Furthermore, these results strongly support CUTS’s ability to measure functional TDP-43 levels under cell stress and other paradigms beyond TDP-43 KD.

**Supplementary Figure 3:**
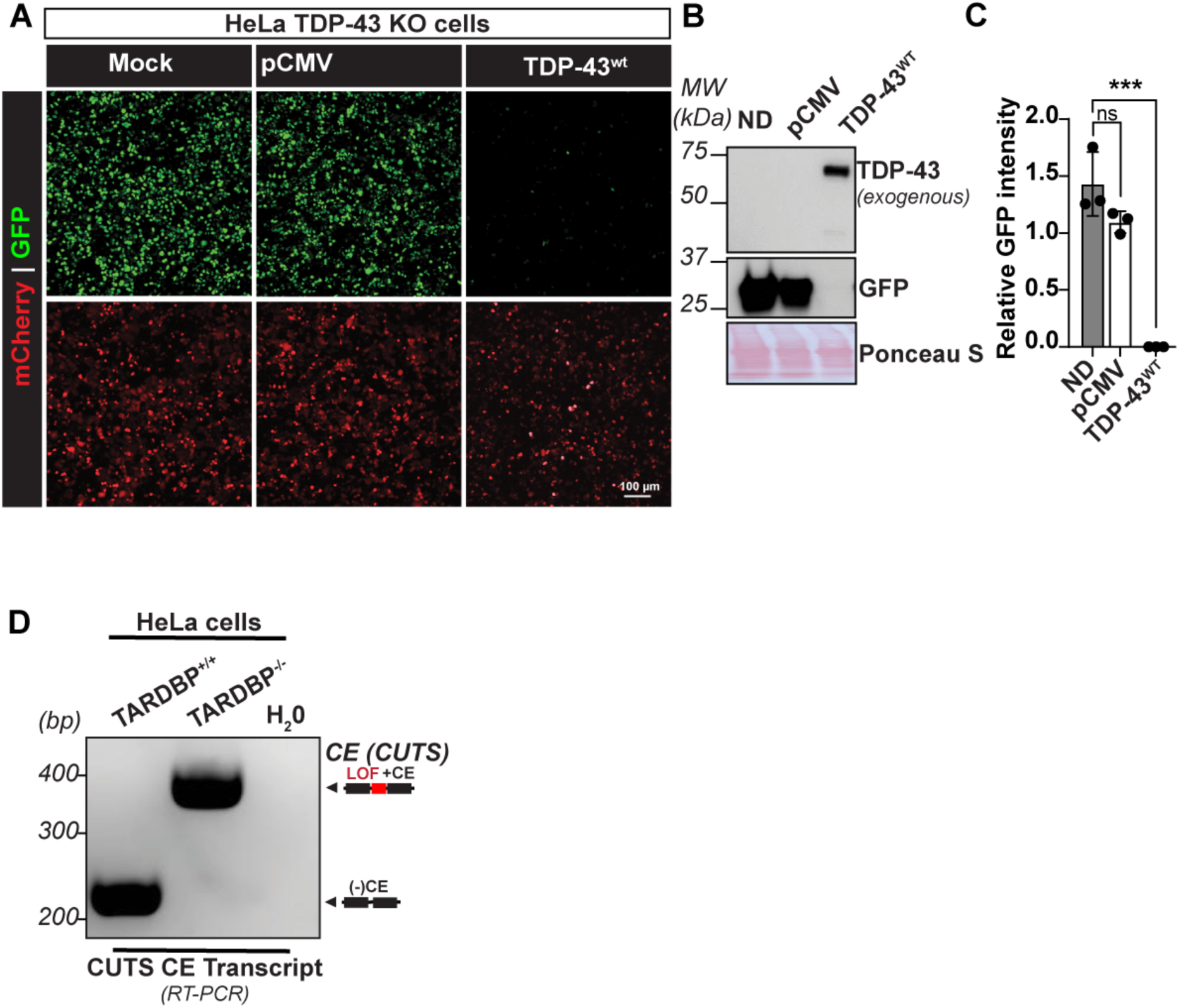
TDP-43^wt^ expression rescues loss-of-function in a HeLa TARDBP Knockout Cell Line. Transient CUTS expression in HeLa TDP-43 KO we induced with doxocycline (1000n/ml) for 24 hours before transfection of pCM backbone or TDP-43^WT^ plasmids. Following transfection, plasmids were expressed for 72 h, followed by live imaging and protein analysis. **(A)** Live-imaging of HeLa TDP-43 KO expressing CUTS in combination with TDP-43^WT^ or pCMV backbone control. **(B)** Representative WB of exogenous and endogenous GFP and TDP-43. Ponceau S is shown as a loading control. **(C)** Relative GFP pixel intensity quantification of the protein bands shown in (B). (**D**)RT-PCR product from CUTS cryptic exon inclusion from wildtype or TDP-43 KO HeLa cell lines using the CUTS-CE primers shown in Figure 2E. Statistical significance was determined by one-way ANOVA and Tukey’s multiple comparison test (*, p < 0.05; **, p < 0.01; ***, p < 0.001; ****, p < 0.0001). Green = GFP; red = mCherry. Scale bar = 100 µm. N = 3 biological replicates.

### CUTS-mediated autoregulated restoration of TDP-43 splicing

Given the growing recognition of the role TDP-43 LOF is believed to play in disease progression, numerous efforts have been committed to developing rescue methods aimed at re-delivering TDP-43 or other gene payloads to restore its physiological splicing function (Baughn et al., 2023; Mehta et al., 2023; Sun et al., 2017). However, a significant challenge in LOF therapies lies in maintaining precise TDP-43 levels within neurons, as even slight overexpression can lead to GOF toxicity (Johnson et al., 2009; Park et al., 2017; Yang et al., 2022). Consequently, a generalized TDP-43 gene-replacement therapy without genome integration carries a substantial risk of overexpression toxicity. Therefore, a CE biosensor such as CUTS may be used to control cell- and temporal-specific regulation of a gene payload. To test this, we generated a CUTS-controlled TDP-43 (CUTS-TDP43) transgene (Figure 4A). Considering the ultra-sensitivity to TDP-43 LOF and minimal leakage under physiological TDP-43 levels, CUTS-TDP43 may have the potential to autonomously negatively regulate its expression, ensuring levels will not surpass physiological levels.

**Figure 4:**
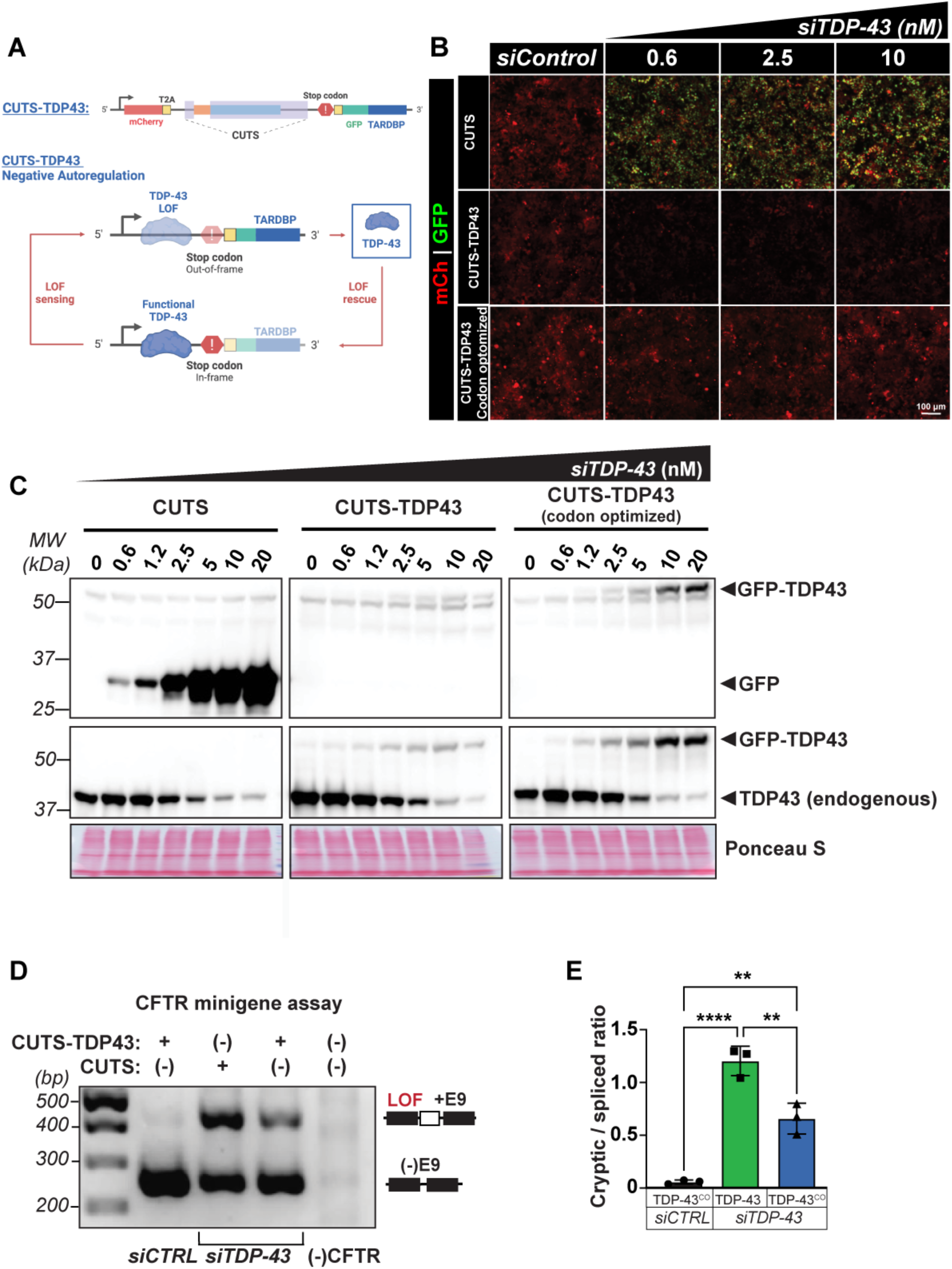
CUTS functions as an autoregulatory controller of TDP-43 expression. **(A)** Schematic of CUTS as an autoregulatory controller of TDP-43 expression (CUTS-TDP43). **(B-C)** TDP-43 siRNA (siTDP-43) dose-response curve in stable polyclonal HEK cells expressing CUTS, CUTS-TDP43, or CUTS-TDP-43^CO^ (codon optimized). The codon-optimized variation allows for continued expression during siTDP-43 treatment. HEK cells expressing the CUTS, CUTS-TDP-43 and CUTS-TDP-43^CO^ were reverse transfected with control siRNA (siCTRL) or siTDP-43 in a dose-response curve (0.6nM-20nM) in a doxycycline (1000ng/ml) supplement media for 72hr. Cells were then used for live imaging or protein analysis. (B) Live imaging of the CUTS variants. (C) Immunoblot assay of GFP and TDP-43. Ponceau S is shown as a loading control. **(D-E)** CFTR minigene assay in stable CUTS or CUTS-TDP-43^CO^ expressing HEK cells. Cells were induced with doxycycline (1000 ng/mL) for 24 h before transfection with the CFTR minigene. Following an additional 24h of expression, cells were transfected with 20nM siCTRL or siTDP-43. Cells were harvested 48 h following siRNA transfection for RNA extraction and RT-PCR analysis. (D) PCR agarose gel of CFTR minigene. (E) PCR analysis of the ratio between the CFTR cryptic exon inclusion and the correctly spliced product from CFTR as shown in (D). Statistical significance was determined by One-Way ANOVA and Tukey Post-hoc (**, p < 0.01; ****, p < 0.0001). Green = GFP; red = mCherry. Scale bar = 100 µm. N=3 biological replicates.

To test this, we created a new polyclonal stable line in HEK293 cells (CUTS-TDP43) by replacing the 3xNLS in the original CUTS cassette with the *TARDBP* ORF fused to a GFP reporter (Figure 4A). However, as our siTDP-43 targets the sequence within the coding region, CUTS-TDP43 was also knocked down upon siRNA transfection, shown by a generalized decreased mCherry signal (Figure 4B). Therefore, we designed a codon-optimized CUTS-TDP43^CO^ that is not targeted by siTDP-43 (Figure 4B). Live imaging analyses showed that CUTS GFP signal demonstrated a steady increase in expression in response to increasing doses of siTDP-43; however, the GFP signal from CUTS-TDP43^CO^ remained undetectable (Figure 4B). WB analysis further demonstrated successful TDP-43 rescue under endogenous TDP-43 KD, as shown by the increasing exogenous TDP-43 observed following decreases in endogenous TDP-43 (Figure 4C). The amount of total TDP-43 appeared to remain consistent throughout the increasing siTDP-43 doses, indicating tight regulation of the rescue parameters. To further confirm whether CUTS-TDP43^CO^ could rescue TDP-43 splicing functionality, we performed a *CFTR* minigene assay (Ayala et al., 2006; Buratti and Baralle, 2001). The expression of CUTS-TDP43^SO^ demonstrated partial, yet significant rescue of cryptic exon 9 splicing in *CFTR* minigene, supporting its controlled efficacy in rescuing splicing LOF (Figure 4D-4E). Taken together, these data indicate that CUTS can autoregulate a TDP-43 payload to physiological levels in response to TDP-43 knockdown.

### Modeling CUTS system in neurons

TDP-43 pathology primarily affects neurons; therefore, we adapted the CUTS system for use in human neuronal models. In the original design, CUTS expression was driven by a Tet-On 3G element under the control of the human phosphoglycerate kinase (hPGK) promoter. While hPGK is generally considered a ubiquitous and stable promoter that drives moderate gene expression and performs well in mitotic cells such as HEK293 and HeLa, several studies have reported reduced activity or silencing of PGK-driven transgenes during neuronal differentiation(Xia et al., 2007). To overcome this limitation, we developed an improved version of CUTS in which the Tet-On 3G element is driven by the human synapsin I (hSYN1) promoter, providing more stable and neuron-specific expression. In this optimized construct, we also replaced mCherry with mScarlet to enhance fluorescence intensity and improve the dynamic range of detection. To test the performance of this updated system, we generated stable human neuroblastoma BE(2)-C cell lines expressing CUTS and examined mScarlett and GFP expression under 50 nM of siTDP-43 treatment in 10-day neuron-differentiated stages (Figure 5A-C). We observed robust and inducible CUTS activation following siTDP-43 treatment in differentiated neurons, as shown by GFP mean intensity in MAP-2 positive cells (Figure 5A-D, S4A). Furthermore, TDP-43 immunostaining demonstrated that cells with reduced TDP-43 protein levels corresponded to GFP-positive cells (Figure 5B).

**Figure 5:**
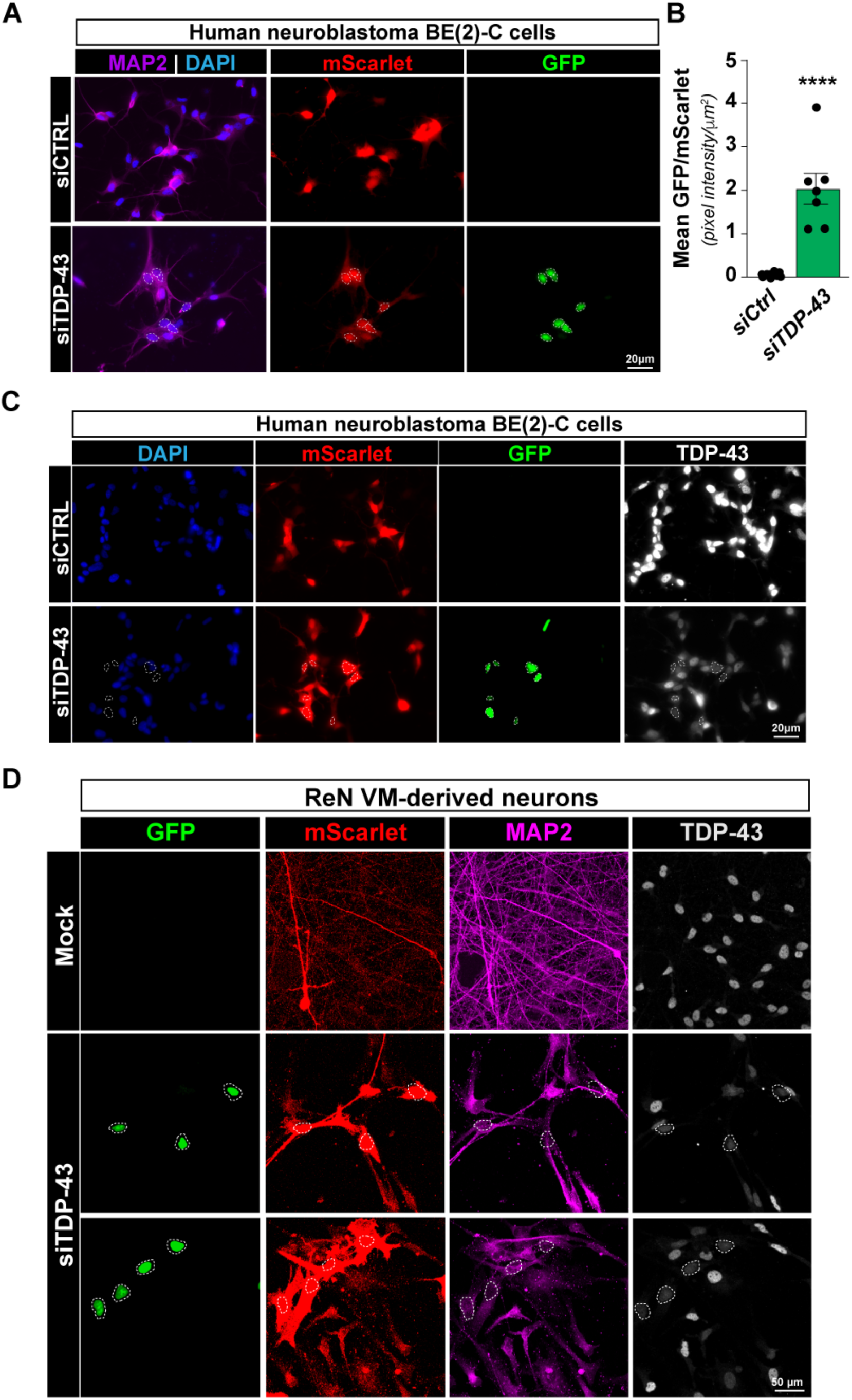
CUTS is activated in human neuronal models in response to TDP-43 LOF. **(A)** Representative immunofluorescence images of MAP-2 (magenta), mScarlet, and GFP from CUTS-expressing BE2C neuron-differentiated cells treated with siCtrl or siTDP-43 (50 nM). Scale bar = 20 μM. **(B)** Mean GFP/mScarlet pixel intensity ratios from (A). **(C)** Representative immunofluorescence images of TDP-43 (greyscale) and mScarlet-GFP signal from CUTS-expressing BE(2)-C neuron-differentiated cells show reduced TDP-43 expression following siRNA treatment. Fluorescence intensity quantification shows a 61.9 +/-9.6 percent knockdown. Scalebar = 20 μM. **(D)** Representative immunofluorescence images of MAP-2 (magenta), TDP-43 (grey), mScarlet, and GFP from CUTS-expressing ReN differentiated neurons with 100 nM siRNA TDP-43. Nuclear GFP signals are indicated with a white dotted circle. Scalebar = 50 μM. Non-parametric Kolmogorov-Smirnov T-test (****, p < 0.0001).

To validate CUTS performance in another human neuronal model, we generated the CUTS human ReN primary neural cells. Stable CUTS-ReN cell lines were generated using a synapsin-driven Tet3G doxycycline-inducible promoter. CUTS-ReN cells were differentiated for three weeks prior to siTDP-43 treatment with increasing concentrations (25–100nM). Live-cell imaging was performed to monitor mScarlet and GFP signals and revealed nuclear GFP in cells with as little as 25 nM TDP-43 siRNA (Figure S4B). Using 100nM siRNA transfection, immunofluorescence analysis of MAP2 and TDP-43 revealed GFP-positive neurons that concurrently showed reduced nuclear TDP-43 levels (Figure 5D; white dotted lines indicate nuclear GFP). Importantly, these GFP-positive cells were negative for the astrocytic marker GFAP (Figure S4C), confirming neuronal specificity of the CUTS response.

**Supplementary Figure 4:**
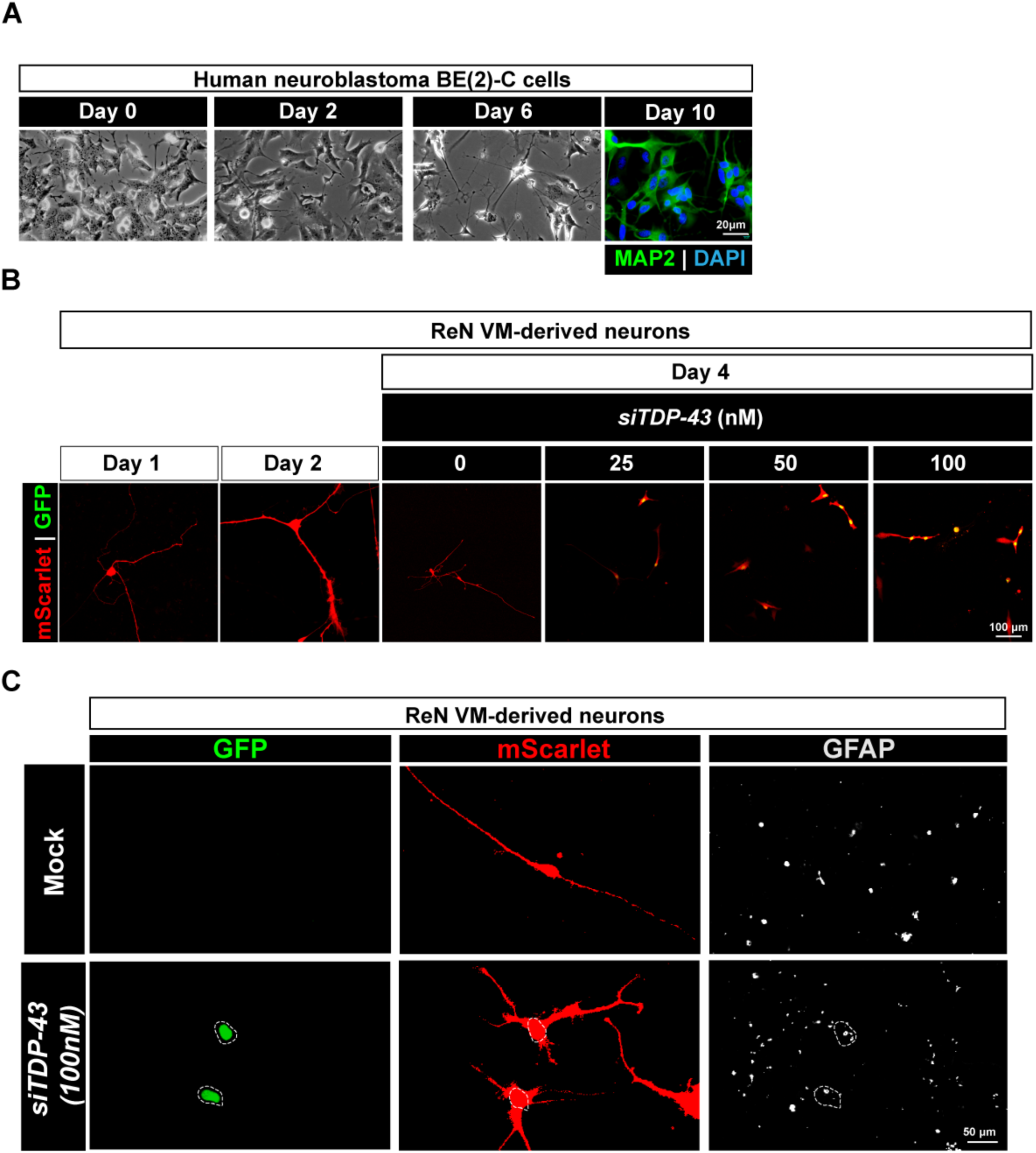
CUTS activity in differentiated BE2C and ReN Cells. **(A)** Representative bright-field images of BE2C differentiation from day 0 to day 6, after 10 days of differentiation, immunofluorescent staining was performed again MAP-2. Nuclei are indicated with a white dotted circle. Scalebar = 20 μM. **(B)** Representative mScarlet and GFP signal from live imagining of differentiated ReN cells expressing CUTS 2 days before siTDP-43 knockdown and after 96 h of siRNA treatment at different siTDP-43 doses. Scalebar = 100 μM. **(C)** Representative immunofluorescent imaging of CUTS system in differentiated ReN VM cells stained for GFAP (Greyscale). Nuclei are indicated with a white dotted circle. Scalebar = 50 mM.

## Discussion

We developed and characterized the CUTS system, a novel approach to detect TDP-43 LOF. The CUTS system utilizes TDP-43-dependent CE events to correlate the level of TDP-43 LOF directly with the expression of a GFP reporting gene. By combining the CFTR-TS and UNC13A-TS, our findings demonstrate that the CUTS system provides an optimal balance of sensitivity and accuracy. This was evidenced by the ability of CUTS to detect modest levels of TDP-43 LOF, as shown by the dose-responsive increase in the expression of the reporter gene, GFP, under various siTDP-43 concentrations (Figure 2). Furthermore, our results suggest that CUTS can effectively discern TDP-43 LOF induced by pathological phase transitions or mislocalization due to TDP-43 mutant or stressor, a critical aspect in the context of neurodegenerative diseases containing TDP-43 pathology, such as ALS and FTLD. The CUTS system’s potential for application in gene-replacement therapies was also highlighted, offering a promising avenue for autoregulated rescue of TDP-43 function, which is critical for avoiding the deleterious effects of TDP-43 overexpression.

This tool findings enables for the significant advancement in the capacity to detect TDP-43 LOF using biosensor assays across diverse experimental settings and through multiple analytical methods. Previously, the *CFTR* minigene assay has been the predominant approach for detecting TDP-43 LOF (Ayala et al., 2006; Buratti and Baralle, 2001; Cohen et al., 2015; Conicella et al., 2020; Jiang et al., 2017; Schmidt et al., 2019). However, this approach is associated with several limitations, all of which are effectively addressed by utilizing the CUTS system. The first advantage of the CUTS system is its ability to detect TDP-43 LOF in real-time through live-imaging analysis. Unlike *CFTR* minigene assays, which typically necessitate endpoint experimental analysis, CUTS facilitates continuous monitoring, eliminating the need for multiple fixed time points. Additionally, CUTS can be seamlessly integrated with various analytical methods, including RT-qPCR (at the RNA level), WB analysis (at the protein level), live imaging (for real-time assessment), and immunofluorescence imaging (to correlate with relevant markers). While not evaluated in this study, it is conceivable that CUTS could be adapted for use with flow cytometry-based techniques, leveraging GFP-positive cells as an output for analysis, as previously demonstrated with a *CFTR*-modified sensor.

In addition to the expanded array of analytical methods offered by CUTS compared to *CFTR* minigene assays, we anticipate that the CUTS system will exhibit superior sensitivity and accuracy. This is supported by the comparison of CUTS with the CFTR-TS or UNC13A-TS cassette (Figure 1), underscoring the potential of CUTS to outperform single minigene-based approaches in TDP-43 LOF detection. Recent work characterizing other CE biosensors indicates that CUTS exhibits comparable or, in some assays, enhanced sensitivity (Wilkins et al., 2024). The expression of the GFP reporter in CUTS achieved up to 118,224-fold increase upon TDP-43 knockdown, compared to *ADNP2* (< 5-fold)(Zhang et al., 2024); TDP-REGv1 (<20-fold); and TDP-REGv2 (<300-fold)(Wilkins et al., 2024). We also show that CUTS can detect ultra-low levels of TDP-43 knockdown (increasing > 7-fold), below the WB or RT-qPCR detection limit. Furthermore, CUTS exhibits a robust log-linear relationship to siRNA doses, making it suitable for quantitative purposes.

The CUTS system outperformed traditional Western blot and transcript-based assays by detecting as little as 1-2% TDP-43 knockdown, which was below the threshold of conventional protein detection methods. This ultra-sensitivity aligns with recent evidence showing that even subtle reductions in TDP-43 can disrupt splicing homeostasis and induce cryptic exon inclusion in physiological targets such as *HDGFL2*, *ARHGAP32*, and *ACBD3* (Brown et al., 2022; Ma et al., 2023; Fratta et al., 2018). The linear correlation between siTDP-43 dose and CUTS reporter output underscores its predictive potential for quantifying graded LOF states. Importantly, the observation that *HDGFL2* cryptic exon inclusion occurs at lower TDP-43 depletion levels supports previous reports identifying it as one of the earliest and most sensitive markers of TDP-43 dysfunction (Humphrey et al., 2020; Seddighi et al., 2023). Thus, CUTS represents a robust and scalable platform to detect and model early TDP-43-dependent splicing alterations.

Due to its success using *in vitro* models, we anticipate that CUTS might similarly be used *in vivo* when coupled with disease models. Integrating the CUTS system with *in vitro* or *in vivo* disease models enables the evaluation of the model’s fidelity in recapitulating TDP-43 LOF phenotypes. Such assessments are crucial for selecting appropriate models that faithfully replicate the desired study context. Furthermore, coupling the CUTS system with TDP-43 models presents a valuable approach for diverse screening studies. For example, CUTS can be leveraged for high-throughput drug screening and CRISPR screening methodologies. Such approaches hold promise for uncovering critical insights into cell-specific disease mechanisms, identifying pivotal disease modifiers, and delineating potential therapeutic genetic targets (Aldewachi et al., 2021; Bock et al., 2022).

A significant advantage of the CUTS system lies in its capacity to deliver precisely regulated gene therapy for rescuing TDP-43 LOF. This study illustrates this capability by placing a functional TDP-43 transcript downstream of the CUTS regulatory elements. The system’s self-regulating ability enhances its safety profile as a gene therapy approach, ensuring gene expression occurs only when necessary and exclusively in cells lacking TDP-43 function. Furthermore, this system can be expanded by substituting the TDP-43 transcript with other genetic modifiers of disease, such as antibodies and PROTACs (Pozzi et al., 2019; Tseng et al., 2023), or genes with established therapeutic potential, including heat shock proteins (HSPs) or heterogeneous nuclear ribonucleoproteins (hnRNPs) (Koike et al., 2023; Lu et al., 2022; Yu et al., 2021). This adaptability holds promise for achieving safe therapeutic outcomes without the need for direct TDP-43 expression.

## Methods

### Generation of plasmids

The CUTS sequence, plasmid, and map were originally generated in this study. The CFTR-TS, UNC13A-TS, CUTS, DNA sequences were designed *in silico* and *de novo* synthesized by Genewiz. These sequences were assembled into Tet3G vector between EcoRI and NotI with NEBuilder HiFi DNA Assembly Master Mix (NEB, E2621L) following the manufacturer’s protocol. For CUTS:hSNY1 construction hSYN1 promoter and mScarlet DNA fragment were generated in GenScript and TRE3G promoter was cut from CUTS plasmid by using ClaI and AfeI. After gel recovery (D4001, ZYMO research) these sequences were assembled into Tet3G vector containing CUTS sequence between PPuMI and XmaI with NEBuilder HiFi DNA Assembly Master Mix (NEB, E2621L) following the manufacturer’s protocol. Transform mix to competent cell (C2987H, NEB) for the following ampicillin selection. Recombined plasmid was extracted by QIAprep Spin Miniprep Kit (27104, QIAGEN) and confirmed by whole-plasmid sequencing. The full DNA sequence for CUTS, CFTR-TS, UNC13A-TS and CUTS:hSNY1 can be found in Table S1.

The CFTR minigene assay plasmid (pTB-CFTR-A455E) was a kind gift from Dr. Yuna Ayala. The exogenous TDP-43 plasmids were constructed in a pCMV backbone by linking a 3xFlag-APEX2 protein (Addgene #164622) (Bonet-Ponce et al., 2020) to TDP-43 coding sequences with WT, cyto, 5FL, or cyto 5FL modifications (Mann et al., 2019).

The codon-optimized *TARDBP* coding sequence (Table S1) was synthesized by IDT and assembled downstream of the GFP sequence of CUTS with NEBuilder HiFi DNA Assembly Master Mix.

All the primers were synthesized by IDT. All plasmids were verified using whole-plasmid sequencing via Oxford Nanopore, provided by Plasmidsaurus.

### Cell culture and transfection

Human Embryonic Kidney 293 (HEK293) cells (female genotype, acquired from the American Type Culture Collection (ATCC)) and HeLa TDP-43 knock-out (KO) cells (a kind gift from Dr. Shawn M Ferguson) (Roczniak-Ferguson and Ferguson, 2019) were cultivated in Dulbecco’s Modified Eagle Medium high glucose, pyruvate (DMEM, Thermo Fisher Scientific, 10-313-039) supplemented with 10% HyClone Bovine Growth Serum (Cytiva HyClon, SH3054103HI) and 1X GlutaMAX (Thermo Fisher Scientific, 10-313-039). Cells were incubated at 37°C in a 5% CO2 atmosphere with high humidity. For transfection assays, cells were plated on collagen-coated coverslips or dishes (50 μg/mL, GIBCO) and transfected with designated DNA quantities using Lipofectamine 3000 (Thermo Scientific, L3000015) following the provider’s protocol. BE(2)-C neuroblastoma cells were maintained in DMEM/F12 (Gibco; Dulbecco’s Modified Eagle Medium/Nutrient Mixture F-12 with GlutaMAX™) supplemented with 10% FBS and cultured until reaching 80–90% confluence. Neuronal differentiation was induced with 10 µM retinoic acid and 50 ng/mL BDNF for 10 days, as described previously (Targett et al., 2024). ReN VM cells were maintained in proliferation medium consisting of DMEM/F12, B27 supplement, hEGF (20 ng/mL), bFGF (20 ng/mL), and heparin (2 ng/mL) on Matrigel-coated plates. Once cultures reached ∼90% confluence, the medium was replaced with differentiation medium (DMEM/F12, B27, and 2 ng/mL heparin) and cells were differentiated for 3 weeks. Full medium changes were performed every other day during the first week, followed by half-medium changes every 4 days thereafter.

### Stable cell line generation via PiggyBac transposition

For stable cell line creation, HEK293 or BE(2)-C neuroblastoma cells were pre-plated on 6-well plates and transfected at approximately 70% confluence with 2.5 μg of PiggyBac plasmids encoding CUTS, CFTR-TS, UNC13A-TS, and CUTS-TDP43 for HEKs cells or CUTS:hSNY1 for BE(2)-C alongside with 0.5 μg of the Super PiggyBac Transposase Expressing plasmid (PB200PA-1) using Lipofectamine 3000, according to the manufacturer’s guidelines. A transfection control without transposase was included. Following a 48-hour post-transfection period, cells were selected with puromycin (Sigma, P8833) at 5 μg/mL, with media changes every two days. Selection resulted in control cell death within approximately 5 days, while surviving populations were expanded and maintained in reduced puromycin concentrations (2.5 μg/mL) to establish stable lines. Expression of the transgenes was confirmed by immunofluorescence staining and Western blot analysis. ReN VM neuronal precursor cells were electroporated using the 4D-Nucleofector system (Lonza). Cells were subcultured two days before nucleofection to reach ∼80% confluence. At confluence, 7.5 × 10⁵ cells were collected, centrifuged, and resuspended in SF buffer (Lonza) containing 0.4 µg of PiggyBac plasmid encoding a CUTS:hSNY1, which is human synapsin driving the Tet-On 3G transactivator gene to activate CUTS expression upon doxycycline treatment, and 0.1 µg PiggyBac transposase plasmid. Electroporation was performed in 16-well Nucleocuvette strips using the CM-137 program. Following a 48-hour post-transfection period, cells were selected with puromycin (Sigma, P8833) at 1 μg/mL, with media changes every two days. Selected CUTS:hSYN1 ReN VM neuronal polyclonal precursor cells were differentiated for 3 weeks. Expression of the CUTS was confirmed in CUTS:hSNY1 ReN VM neurons by live imaging and immunofluorescent staining following treatment with 1000 ng/ml of Doxycycline for 5 days.

### Cellular stressor treatment

Sodium arsenite (NaAsO₂) was prepared in fresh culture medium to a final concentration of 250 µM and applied to cells by complete medium exchange. For recovery, cells were washed once with fresh medium and subsequently incubated in recovery medium for 5 hours.

### SDS-PAGE and Western blot

For protein analysis, cells were lysed directly on the plate using fresh and pre-chilled Urea-RIPA buffer: 2M fresh urea in 1XRIPA buffer (Boston Bioproducts, BP-115X), supplemented with 1% protease inhibitor cocktail (Sigma, P8340) and sonicated. Protein concentrations were quantified using the Pierce BCA Protein Assay Kit (Thermo Scientific, 23227). Proteins were resolved by SDS-PAGE and transferred to nitrocellulose membranes for WB analysis. Membranes were blocked and probed with primary antibodies: mouse-anti-GFP (Santa Cruz, sc-9996, 1:200), mouse-anti-α-tubulin (Sigma, T5168, 1:1000), rabbit-anti-TDP-43 (Proteintech, 10782-2-AP, 1:2500), and rabbit-anti-mCherry (Cell Signaling, 43590, 1:1000), followed by HRP-conjugated secondary antibodies: donkey-anti-mouse (JacksonImmunoResearch 715035151, 1:5000) or donkey-anti-rabbit (JacksonImmunoResearch 711035152, 1:5000). Detection was achieved using Western Lightning ECL Pro (Revvity, NEL1201001EA) or Supersignal West Femto Maximum Sensitivity Chemiluminescent Substrate (Thermo Scientific, 34095) in an Amersham ImageQuant 800 GxP biomolecular imager system (Amersham, 29653452).

### Fluorescence microscopy

Confocal imaging was performed on a Nikon A1 laser-scanning microscope using either a 60X oil immersion or a 10X/20X objective for live-cell observations. A Tokai HIT stage-top incubator maintained the required environmental conditions. Representative images were chosen from at least two independent experiments with a minimum of three biological replicates each. For live imaging quantification, we measured the mean GFP signal intensity for each experimental condition. The values were then averaged, and the results were used to calculate and plot the fold change. For immunofluorescent imaging, we first created maximum intensity projection images. We then applied masks to the GFP, mCherry/mScarlet, and Hoechst signals. By overlapping the GFP and mCherry/mScarlet signals, we identified the number of GFP-positive cells. Similarly, by overlapping the mCherry/mScarlet signal with the Hoechst mask, we identified the CUTS-expressing cells. We then calculated a ratio of GFP-positive cells to CUTS-expressing cells, plotted as a percentage of GFP-positive cells using the Nikon NIS software. Neuron-differentiated BE(2)-C cells were stained for MAP2 (Millipore Cat# A5622, 1:500), TDP-43 (Proteintech, 10782-2-AP, 1:500) and Hoechst (1:10,000) and imaged using an Olympus IX71 inverted microscope equipped with 40X objective, a DP80 camera, and Olympus cellSens software. Fluorescence intensity ratios were calculated across multiple image fields using ImageJ FIJI.

### siRNA

All siRNA treatments were performed as described using RNAiMAX (Thermo Scientific, 13778150) per manufacturers protocols. All siRNA experiments in CUTS:hSNY1 ReN VM neurons were performed with RNAiMAX reagent and siRNA 48 h after Doxycycline treatment (1000 ng/ml). Quantification occurred and quantified after 120 hr post RNAi treatment (168 hr post Doxycycline treatment). Reverse transfections of siRNA were conducted using RNAiMAX reagent adhering to the supplier’s protocol. To knockdown TDP-43, the following siRNAs were used: ON-TARGETplus SMARTpool siRNA against *TARDBP* (Dharmaco, L-012394-00-0005) and siGENOME non-Targeting siRNA for control (Dharmaco, D-001206-13-05).

### RNA extraction, RT-PCR, and qPCR

The RT-PCR or RT-qPCR was conducted with cDNA diluted 10-fold. For RT-PCR assay, CFTR cryptic exon region was amplified with the following primer pair: P690-F (5’-CAACTTCAAGCTCCTAAGCCACTGCCTGC) and P691-R(5’TAGGATCCGGTCACCAGGAAGTTGGTTAAATCA). CUTS’ cryptic exon region was amplified with the following primer pair: CUTS-CE-F (5’-ATCCCGGCCCTGGATCCG) and CUTS-CE-R (5’-GTCAGCTTGCCGTAGGTGGC). Endogenous cryptic exons including *LRP8*, *EPB41L4A*, *ARHGAP32*, *ACBD3* and *HDGFL2* were amplified with the following primer pair: LRP8-CE-F(5’-GGACGAGTTCCAGTGTGGG) and LRP8-CE-R(5’ GAAATCTGCGGGGACCCT); EPB41L4A-CE F (5’AGTCACCTTACAAAACAGAAGTCACA) and EPB41L4A-CE-R(5’-ACATATGCACACACACTCTCACA); ARHGAP32-CE-F (5’GGAGGAAGCATTTTGGAGGTTTCTA) and ARHGAP32-CE-R (5’-CAGCCAAATGCACAGCGAAT); ACBD3-CE-F (5’-ACAGTATCCAGGGAACTACGAA) and ACBD3-CE-R (5’-GCTTCCAGAGAAAGTACCTGTTGC); HDGFL2-CE-F (5’-AAGACGCCTGCGCTAAAGAT) and HDGFL2-CE-R(5’-GCTTCCCTCCCTTCTGATGC). TDP-43 and 18s were amplified using the following primers *RPS18*_FW (5’-GCAGAATCCACGCCAGTACA) and RPS18_REV (5’-TTC ACGGAGCTTGTTGTCCA); *huTDP-43*_FW (5’-TCATCCCCAAGCCATTCAGG) and REV (5’-TGC TTA GGT TCG GCA TTG GA). PCR products were separated by agarose gel electrophoresis, and the bands were visualized with Amersham ImageQuant 800 GxP biomolecular imager system.

For RT-qPCR assay, SsoAdvanced™ Universal SYBR Green Supermix (Biorad, 1725272) was used following the supplier’s protocol on a CFX96 Touch Real-Time PCR Detection System (Biorad). Three technical replicates were included for each sample with the following program: 95°C for 30 s, 40 cycles of 95°C for 15 s and 60°C for 20 s. CUTS-CE-F (5’-ATCCCGGCCCTGGATCCG) and CUTS-CE-R (5’-GTCAGCTTGCCGTAGGTGGC) were used to quantify normal CUTS transcript. CUTS-J-F (5’-TCCGGCGAGGGATTTGGG) and CUTS-J-R (5’-CCCCACCTAGACCCATCTCTCC) were primers targeting the cryptic exon junctions to quantify cryptic exon-specific CUTS transcripts. Relative quantification of CUTS cryptic exon was determined by the ΔCt value of CUTS-J normalized to CUTS-CE.

### Statistical analysis

Statistical significance was evaluated using GraphPad Prism 9 software, and specific tests used for each experiment are outlined in the respective figure legends.

## Authorship contributions

LX, JM, and CAB share equal contributions and authorship. LX and BTH conceptualized the study. LX, JM, and CAB were responsible for data curation, data analysis, conceptualizing and performing the described methodologies, compiling and generating data, and writing the original draft. JX assisted with data analyses and experimental optimization. SB and CTC contributed to experimental design and analysis of the neuronal BE(2)-C studies and edited the manuscript, with SB additionally constructing the stable BE(2)-C lines, conducting the experiments and collecting data. CJD conceptualized the approach, funded the research, and prepared the final manuscript.

## Acknowledgments

We thank Dr. Yuna Ayala, Ph.D. (St. Louis University School of Medicine), for kindly providing the CFTR minigene plasmids. We thank Dr. Shawn M Ferguson, Ph.D. (Yale University School of Medicine) for kindly providing the TARDBP^-/-^ HeLa cell lines used in this study. We thank Olivia R. Shapiro and Jocelyn C. Mauna (Donnelly Lab, University of Pittsburgh School of Medicine) for their kind assistance and helpful discussions. This work was supported by funds to C.J.D. by the LiveLikeLou Fund at the Pittsburgh Foundation and grants from NIH (R01NS105756, R01NS127187) and R21AG075814 to C.T.C.

---

**Supplementary Table 1.**
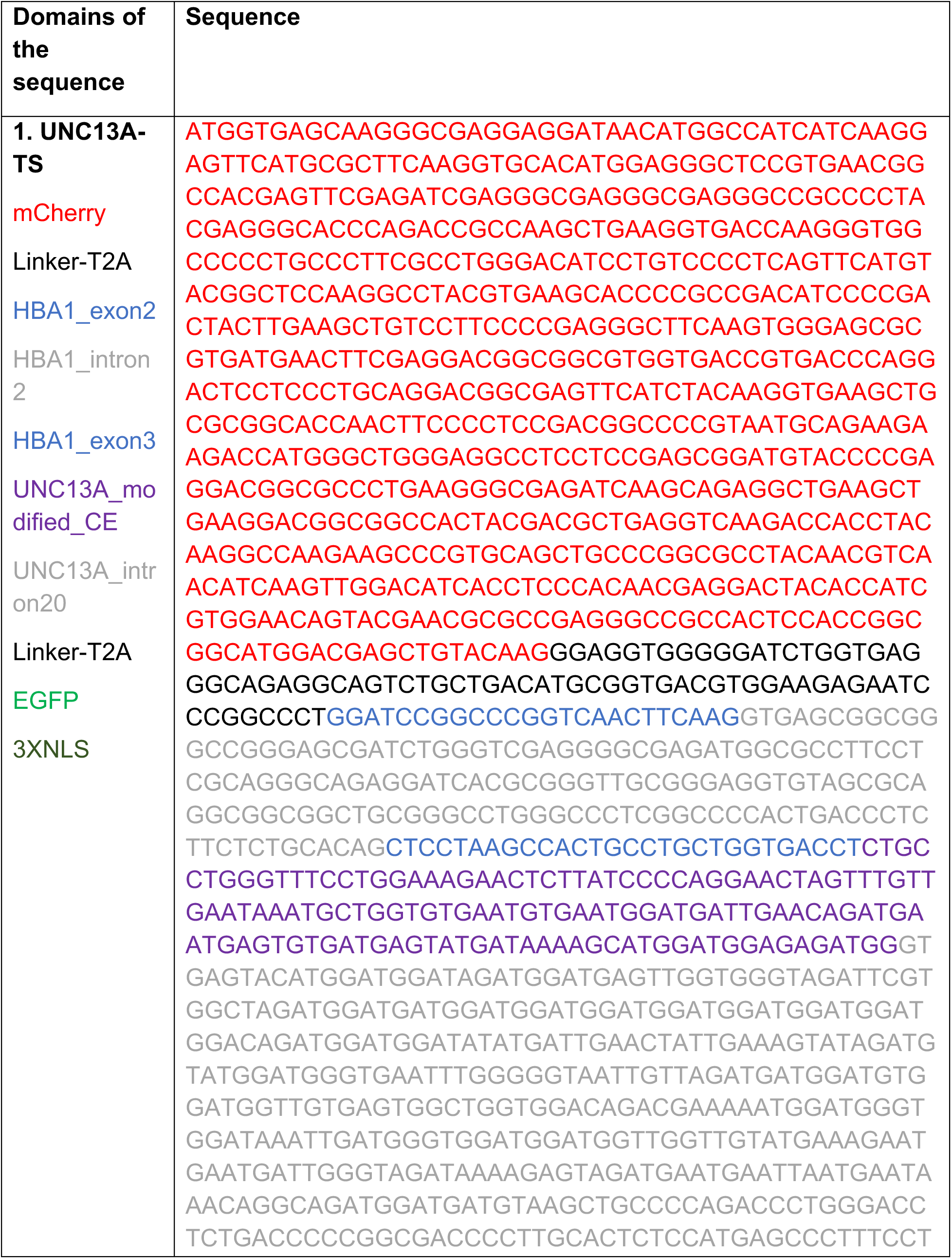
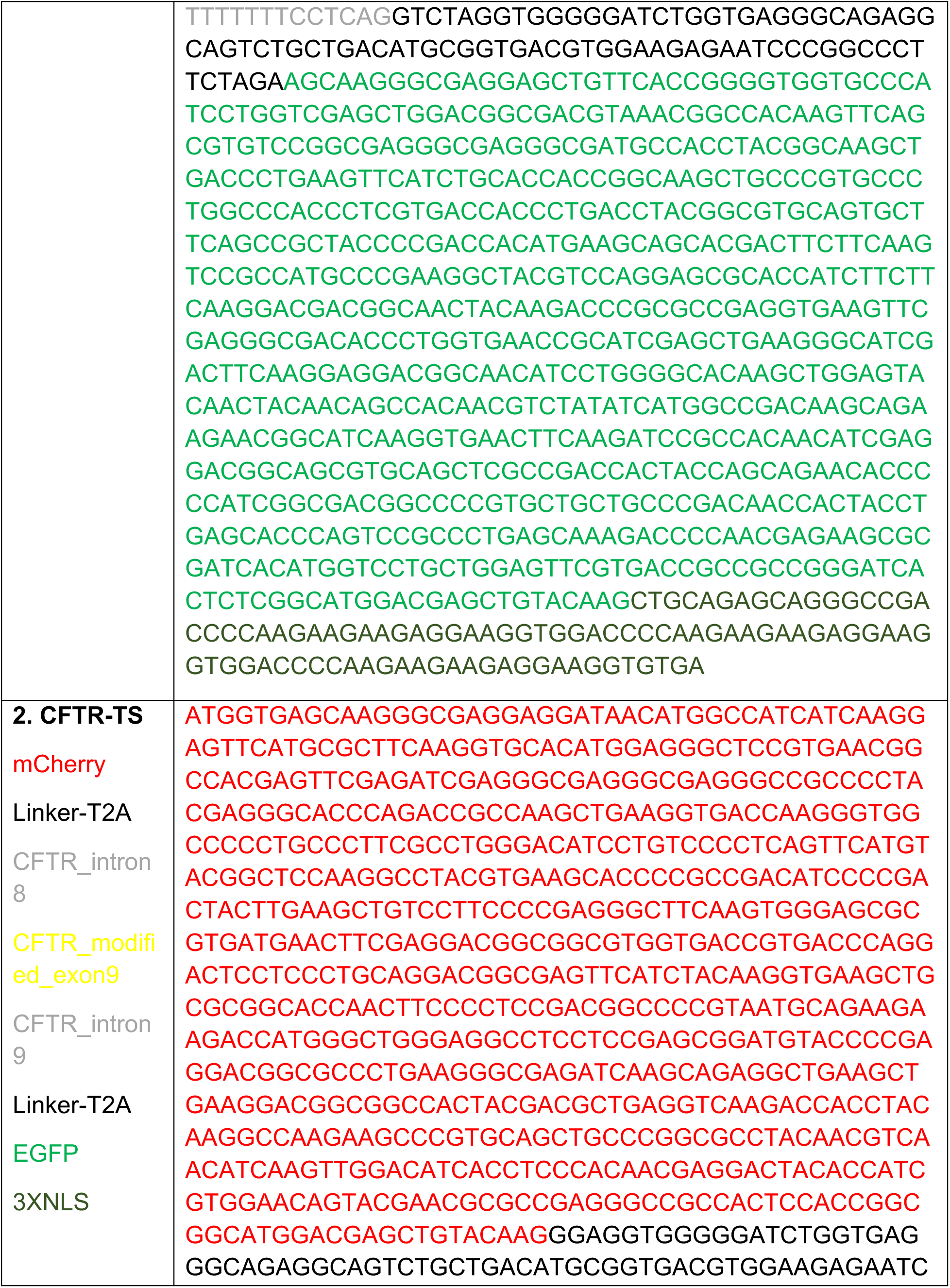
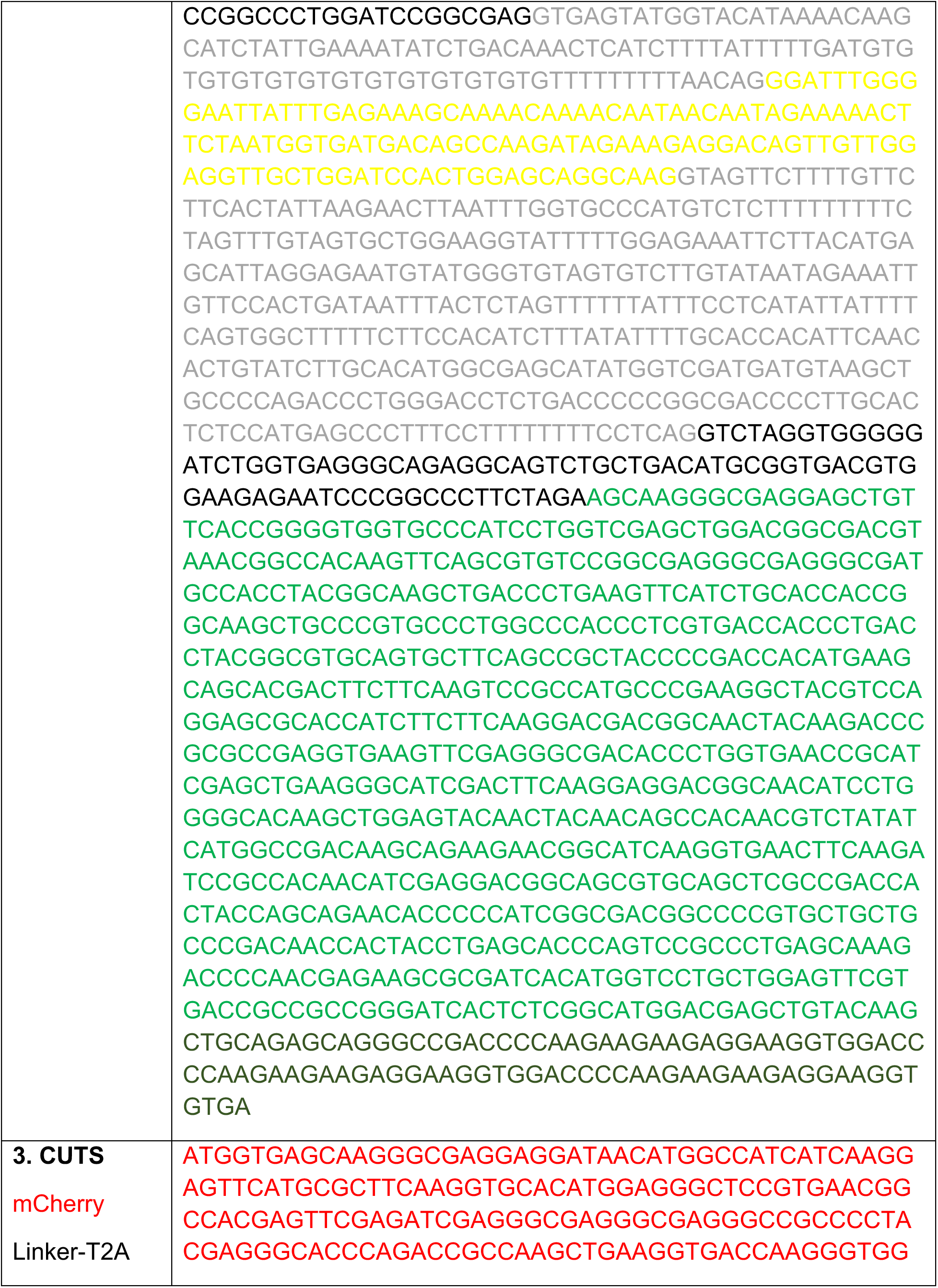
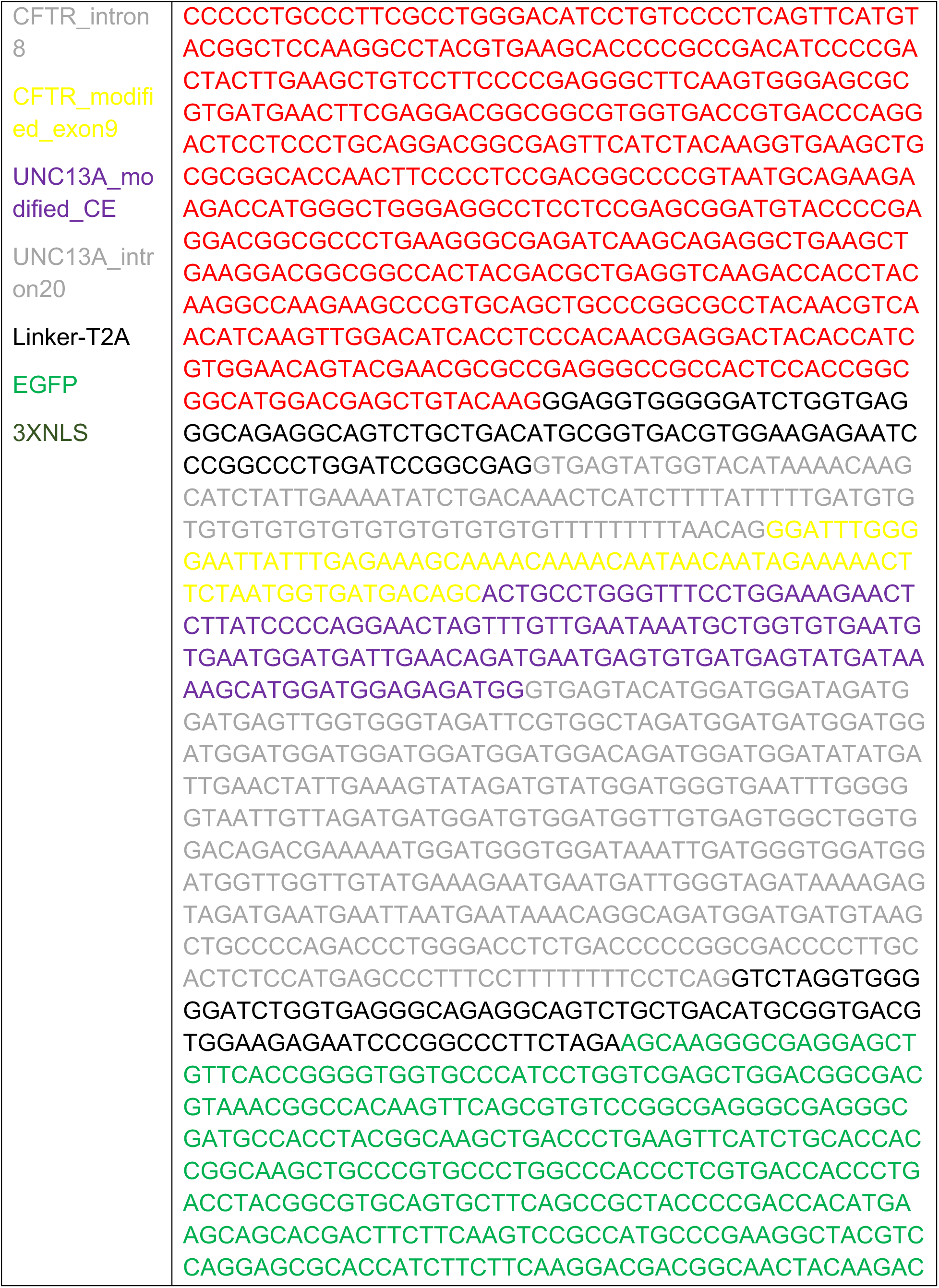
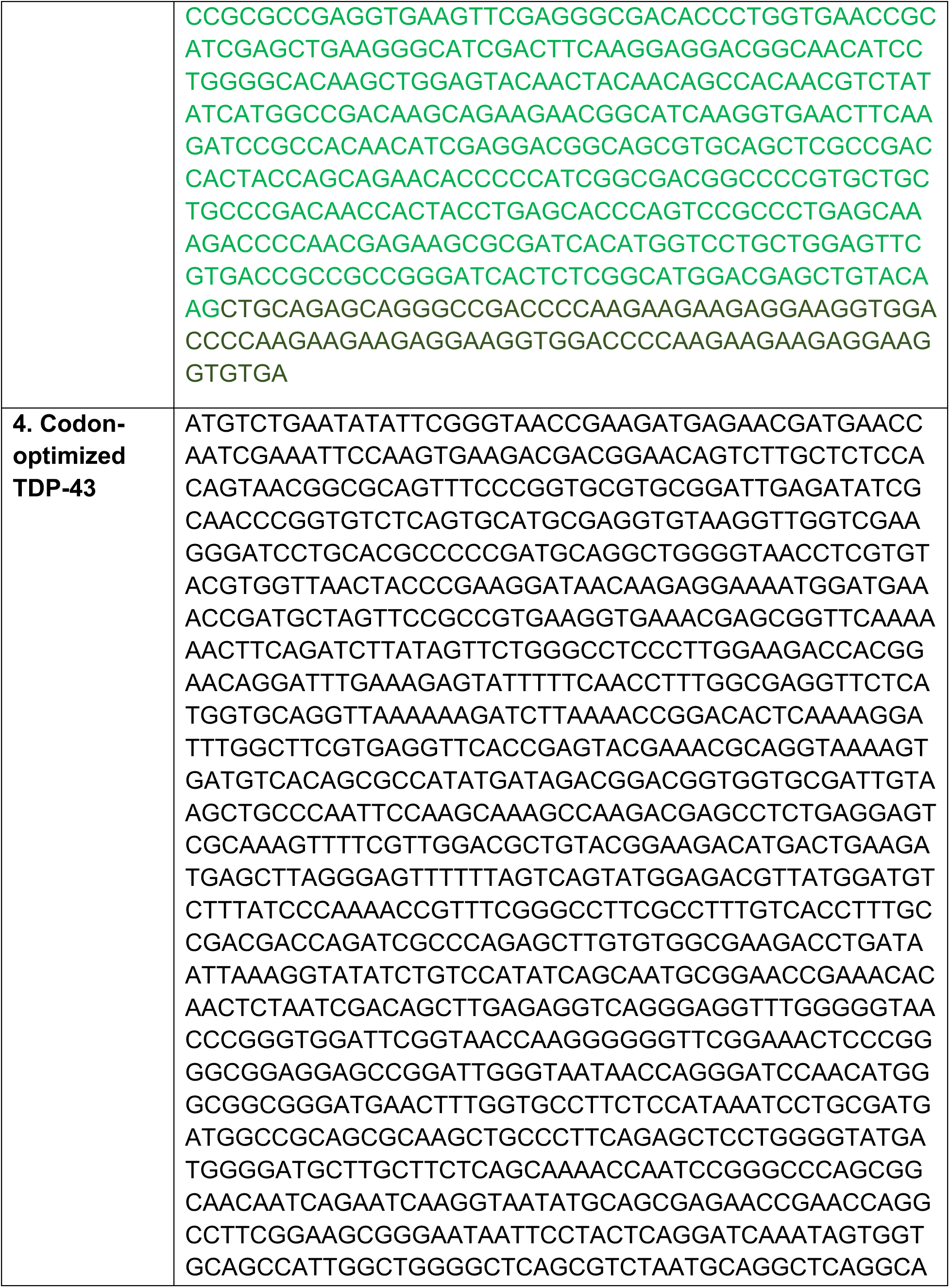
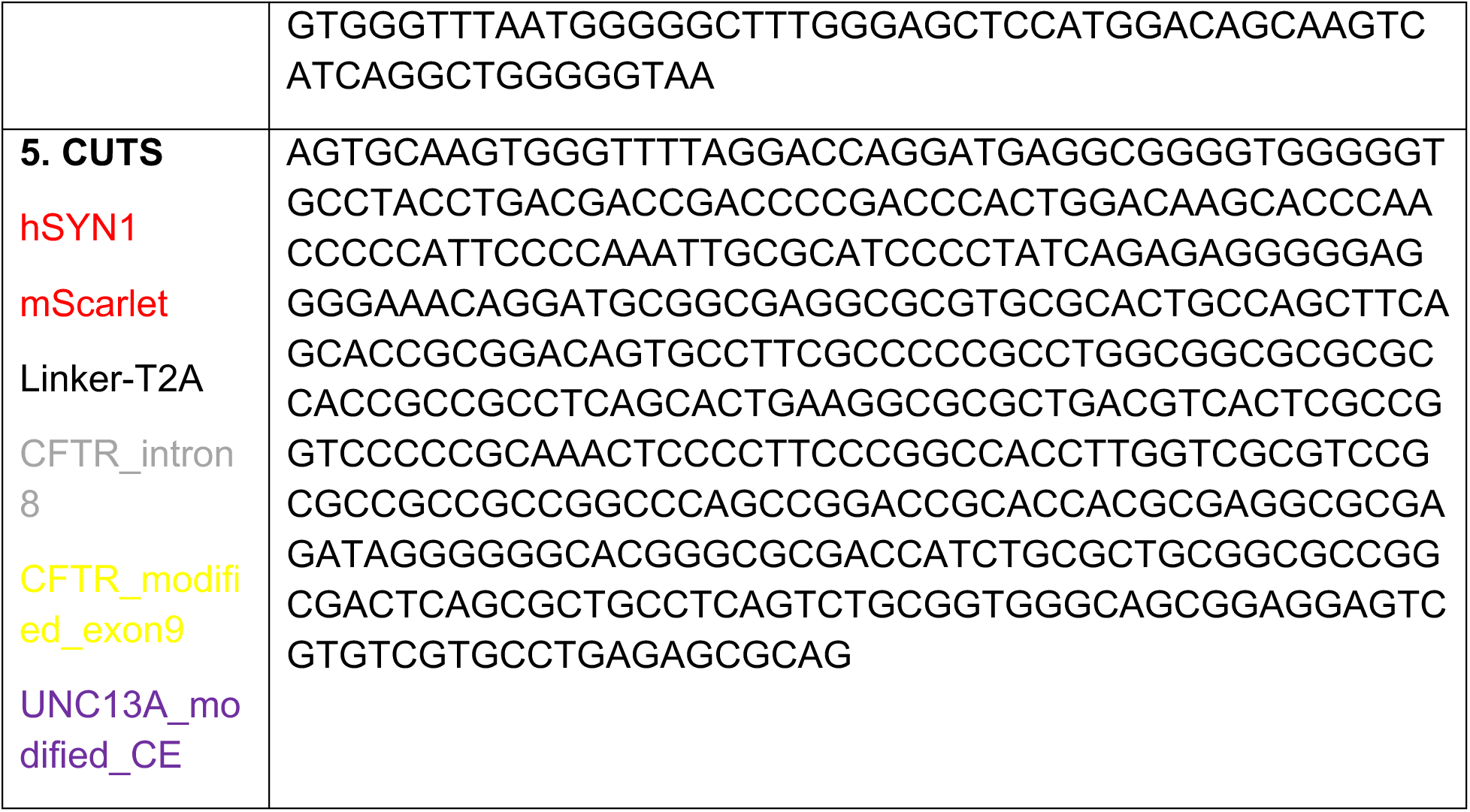
Information of DNA sequence.

